# Comprehensive Mapping of the Virus and Host Factors that Guide the Paths of HIV-1 Escape from a Therapeutic

**DOI:** 10.64898/2025.12.27.696684

**Authors:** Aaron N Gillman, Cassian M Birler, Rohith Rao Vujjini, Mohammad Fili, Samuel A McCarthy-Potter, Wilson Chen, Madeline M Broghammer, Guiping Hu, Margaret J Gartland, Manyu Prakash, Annika Helverson, Grant Brown, Hillel Haim

## Abstract

HIV-1 resistance to therapeutics can emerge through diverse mutational routes, yet the determinants guiding pathway selection *in vivo* remain unclear. Through comprehensive screening, we identified 18 mutations in the HIV-1 Env protein that enhance resistance to the FDA-approved small-molecule therapeutic temsavir. We then examined their occurrence in HIV-infected individuals who developed resistance on therapy. Only a subset of the resistance-enhancing mutations emerged *in vivo*. On-treatment mutation frequencies correlated with their spontaneous emergence rates in temsavir-untreated individuals, and were governed by two parameters: **(i)** Probability of mutation appearance, determined by number and type of nucleotide changes required, and **(ii)** Probability of mutation persistence, determined by Env functional and immune fitness. Notably, non-neutralizing antibodies commonly-elicited in HIV-infected individuals restricted emergence of multiple resistant forms, driving convergence to a narrow set of escape routes. These findings establish a quantitative framework for predicting therapeutic resistance and reveal how host-immunity constrains viral evolution during treatment.

## INTRODUCTION

Human immunodeficiency virus type 1 (HIV-1) has a remarkable ability to adapt to the host environment. Due to the low fidelity of the viral replication machinery (1, 2), nucleotide substitutions are introduced randomly across the genome, allowing the virus to sample new variants of its proteins and regulatory elements (3, 4). Such changes facilitate the broadening of cell tropism (5, 6), evasion of the host immune responses (7, 8), and development of resistance to antiviral therapeutics (9). Clinical resistance can emerge by *de novo* mutations that occur during treatment or by resurfacing of archived forms from latent cell reservoirs (10). Given that therapeutics dock to their protein targets via interaction with multiple residues, several mutational pathways are available for the virus to evade recognition. However, as most antiretrovirals target sites associated with protein function, resistance-enhancing mutations often compromise virus fitness, limiting the ability of these variants to persist in the host. In such cases, adaptive mutations that increase fitness yet retain the resistance phenotype can facilitate persistence (11-13).

While functional fitness and inhibitor resistance are clearly traits essential for virus replication, the principles that govern selection of escape paths from therapeutics remain incompletely understood. First, there is a need for a more quantitative understanding of the contributions of these two traits. For example, is a modestly-resistant but highly-fit variant more likely to persist than a highly-resistant but poorly-fit form, and what thresholds define these distinctions? Second, to what extent do factors other than functional fitness and inhibitor resistance contribute to appearance and persistence of the new variants? For example, amino acid substitutions that require a single nucleotide change are more likely than those requiring two changes. However, the likelihoods for all nucleotide changes, at least *in vitro*, are not identical, with transitions exhibiting higher rates than transversions (14-16). Whether such preferences influence escape pathway selection *in vivo* is unknown. Third, the extent to which we can identify all possible mutational paths that impart resistance to a therapeutic is still unclear. Indeed, HIV-1 proteins, and particularly the envelope glycoproteins (Envs), exhibit considerable within- and between-host variability (17, 18). Such variability results in strain-specific conformations and allosteric effects. Consequently, some mutations may increase resistance or reduce fitness of some isolates but exert limited effects on others. Context-dependence of mutation effects may limit our ability to estimate the resistance of HIV-1 isolates from sequence data, reducing the potential for personalized antiviral therapies. Context-specific effects may also hinder the generalization of *in vitro* findings to the diverse viruses that circulate in the population.

To address these questions, we investigated the *in vivo* escape paths of HIV-1 from the Env-targeting inhibitor temsavir (TMR, BMS-626529), the active metabolite of the FDA-approved prodrug fostemsavir (19, 20). TMR effectively reduces viral loads in HIV-infected individuals (21-23). In addition, it has been shown to reduce the proinflammatory effects of soluble gp120 and to prevent killing of uninfected CD4-positive cells via effector-mediated mechanisms (24-26). We first conducted a comprehensive effort to identify the mutations that increase HIV-1 resistance to TMR *in vitro*. Then, we examined their representation in participants of the BRIGHTE clinical trial who developed resistance to this agent during treatment (21, 22, 27). Interestingly, only a subset of the resistance-enhancing mutations (**REMs**) identified *in vitro* emerged on treatment. REM emergence frequencies correlated strongly with their spontaneous emergence rates in the fostemsavir-untreated population. *In vitro* analysis of the functional and antigenic properties of all REMs as well as *in silico* modeling of their substitution likelihoods revealed that their rates of emergence *in vivo* were explained well by these parameters. Remarkably, non-neutralizing antibodies against Env, which are commonly produced in HIV-infected individuals, appeared to restrict the range of resistance mutations that emerged *in vivo*. Together, our findings establish a quantitative framework for understanding the emergence of resistance to antiviral therapeutics and inform strategies to predict the rate and paths of escape in each individual.

## RESULTS

### Mutations in the core epitope of TMR account for most, but not all, resistance to TMR

The small-molecule inhibitor TMR is generated by hydrolysis of the orally administered prodrug fostemsavir in the gut (**Fig 1A**) (28-30). TMR targets the CD4-binding pocket of Env and stabilizes the trimer in a CD4-unbound state, preventing activation of the entry cascade by the CD4 receptor (**Fig 1B**) (31). The main contact sites for TMR are the side chains of Env positions 375, 426 and 434, which cradle this molecule in the pocket (27, 32-36). Mutations at position 475 have also been shown to impact resistance (36-39). The consensus amino acid motif at these four sites for TMR-sensitive strains is Ser at position 375 and Met at positions 426, 434 and 475, collectively designated herein the **SM^3^ motif**. Mutations at these sites reduce virus sensitivity to TMR *in vitro* and were associated with clinical resistance in treated individuals (22, 27, 32). Several trials were conducted to test the efficacy of fostemsavir and predecessor compounds. The largest trial, BRIGHTE, enrolled 371 HIV-positive participants, who were treated for up to five years (see trial design in **Fig S1A**) (21-23, 26, 27, 40). Plasma samples were collected from the participants before and during treatment and analyzed by the Monogram Biosciences PhenoSense GT test (**Fig S1B**). The assay is based on amplification of *env* from plasma virus, and bulk cloning of the amplicons into an expression vector (33, 41). The library of plasmids from each sample is then sequenced and used to generate a library of pseudoviruses that is tested for resistance to TMR *in vitro*, as measured by the concentration required to achieve a 50% reduction of infection (IC_50_). The trial thus provided genomic and phenotypic data that we used to establish the escape paths of the virus *in vivo* (see **Data File S1**).

**Figure 1.**
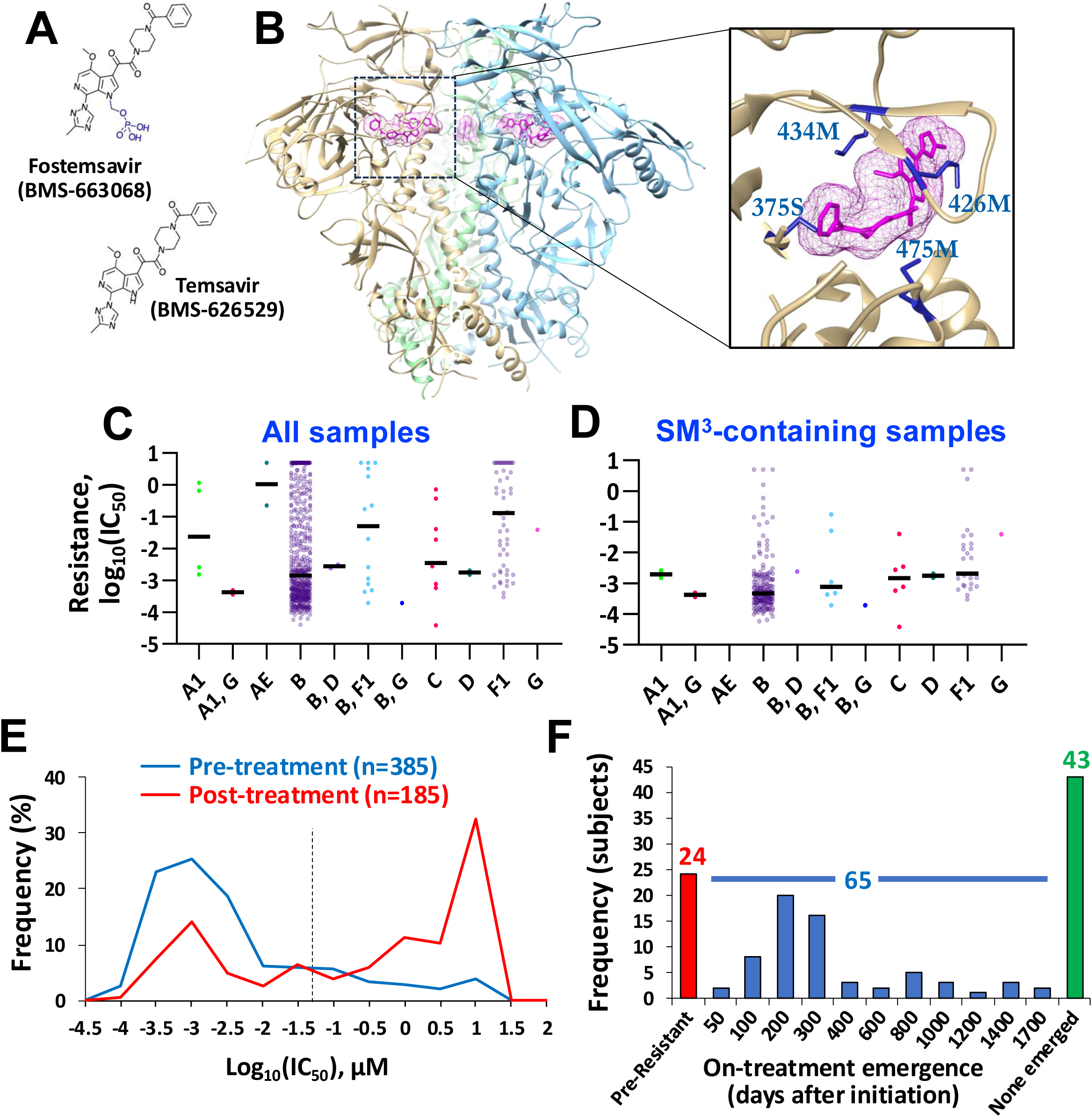
TMR resistance and escape in the BRIGHTE clinical trial. **(A)** Structure of fostemsavir and the active metabolite TMR. **(B)** Cryo-electron microscopy model of the Env trimer bound to TMR (in magenta, PDB ID 8TTW). The inset shows the CD4-binding pocket with the SM^3^ sites labeled. **(C,D)** Resistance values of all 570 samples from BRIGHTE trial participants and of the subset of samples that contain the TMR-sensitive SM^3^ motif. Samples are grouped by their inferred clade associations. **(E)** Distribution of IC_50_ values for samples collected before and after fostemsavir treatment. The threshold used to define resistance (50 nM) is shown by a dotted line. **(F)** TMR resistance outcomes in the 132 BRIGHTE trial participants for whom genotype and IC_50_ data were available for samples collected both before and after treatment.

Sequence and TMR IC_50_ data were available for 570 plasma samples from 360 BRIGHTE participants. We first examined the distribution of IC_50_ values based on their HIV-1 subtype associations (see phylogenetic tree in **Fig S2A** and **Data File S2**). Of the 360 samples, 83.3% were identified as clade B, 3.5% as clade B recombinants, and 9.5% as clade F1. Considerable variability was observed in IC_50_ values within and between the clades (**Fig 1C**). This variability was significantly reduced in the subgroup of samples that contained the SM^3^ motif (**Fig 1D)**. Nevertheless, some SM^3^-containing samples still exhibited high IC_50_ values, suggesting that additional sites impact resistance. We also performed the above analysis for a previously published panel of 208 Envs (designated herein the **Single-Env Dataset**) that were tested for their resistance to TMR (38). A similar decrease in the intra- and inter-clade variability in IC_50_ values was observed for the SM^3^-containing Envs of this panel (**Fig S2B** and **Data File S3**). This finding suggested that sensitivity to TMR involves only modest “clade context” effects and that positions outside the SM^3^ motif may contribute to resistance.

To establish a threshold that distinguishes between sensitive and resistant samples, we examined the distribution of IC_50_ values for the BRIGHTE dataset. A right-skewed distribution was observed for the pre-treatment samples and a bimodal one for the post-treatment samples (**Fig 1E**). Based on these data, we selected an IC_50_ threshold of 50 nM TMR, which represents the 85^th^percentile for the pre-treatment samples. This threshold was then used to define changes in resistance status during treatment. Among the 360 participants, only 132 had sequence and IC_50_ data for both pre- and post-treatment time points (**Fig 1F**). Of these, 24 were resistant to TMR before treatment, 43 remained sensitive to TMR throughout treatment, and 65 gained resistance on treatment (with a median time of 218 days). For this latter group, designated herein the **escape group**, we sought to identify and characterize the mutational paths used by the virus to gain resistance.

### A combined approach to identify Env mutations suspected of increasing HIV-1 resistance to TMR

To identify the escape paths from TMR in the BRIGHTE participants and compare them with all possible paths available to the virus, we pursued the approach described in **Fig 2A**. First, we used four strategies to identify Env mutations suspected of increasing resistance to TMR. We then tested them *in vitro* for their effects on TMR resistance and Env function using a pseudovirus infection assay. Finally, for the resistance-enhancing mutations identified, we examined their emergence frequencies after treatment in the escape group, to determine if the observed frequencies can be explained by their effects on fitness and/or TMR resistance.

**Figure 2.**
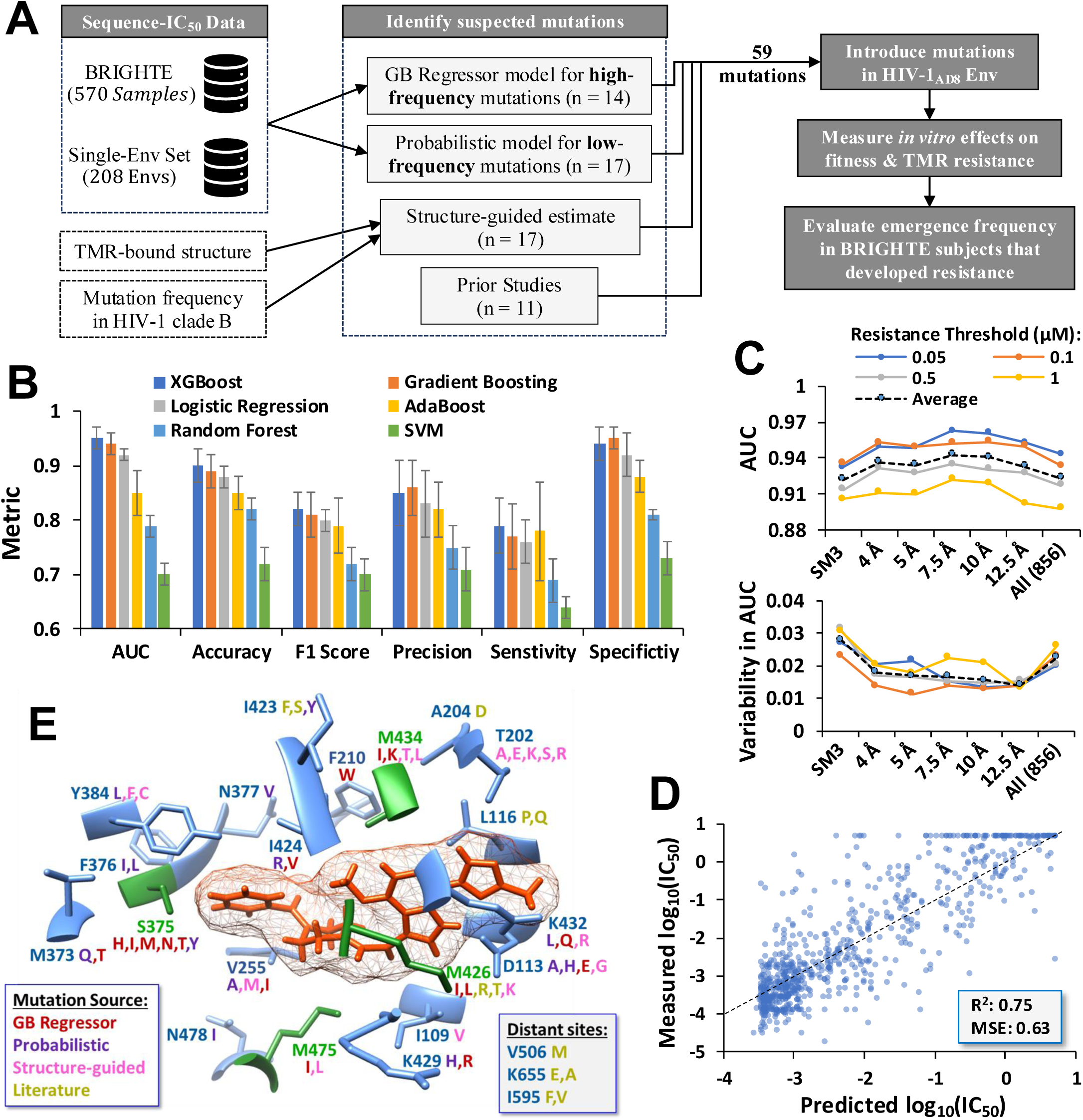
Identification of mutations suspected of increasing HIV-1 resistance to TMR. **(A)** Our approach to identify the mutations. The number of mutations identified by each approach is shown. **(B)** Performance of different algorithms to predict TMR resistance by sequence. Amino acid sequence at the 856 positions of Env in the 570 BRIGHTE trial samples was used as input. Average metrics for five-fold cross-validation are shown. Error bars, standard deviation (SD). **(C)** Performance of the XGBoost algorithm to predict TMR resistance in the BRIGHTE samples by Env sequence. As input, we used amino acids at the four SM^3^ positions, all 856 positions, or positions within the indicated distances from TMR on the TMR-bound structure of Env. Performance was tested using the indicated IC_50_ thresholds to define resistance. Average AUC values and their variability (SD) across the five folds is shown. **(D)** Performance of a Gradient Boosting Regressor model to predict resistance of 778 samples from the BRIGHTE and Single-Env datasets by sequence. MSE, mean squared error. **(E)** The 59 Env mutations suspected of increasing resistance to TMR identified by the four approaches shown in panel

To determine if the genotype-phenotype datasets from the BRIGHTE trial are sufficiently informative to identify mutations that increase resistance to TMR, we tested different machine learning models (see details in the **Methods Section**). As input for the models, we used the amino acids at all 856 positions of Env according to the HXBc2 numbering system (42). The outcome to be predicted was the presence of resistance (IC_50_ value greater than 50 nM). The XGBoost and Gradient Boosting classifier algorithms (43) achieved the highest performance across all key metrics (**Fig 2B** and **Fig S3**), and were thus chosen as the foundations for subsequent models. We then evaluated the optimal number of Env positions to include as input, which were selected based on their minimal distance (in Ångstroms, Å) from the TMR molecule using coordinates of the TMR-liganded Env (PDB ID 5U7O) (38). Positions within 4Å, 5Å, 7.5Å, 10Å or 12.5Å from any TMR atom were tested (see positions in **Table S1**). In addition, we tested the four sites of the SM^3^ motif, and all 856 positions of Env. **Fig 2C** shows the area under the curve (AUC) metric for these tests, which describes performance of the model to distinguish between resistant and sensitive samples by sequence, whereby a value of 1 indicates perfect discrimination and 0.5 corresponds to random classification. AUC values greater than 0.9 were observed for most sets, peaking at 0.96 for the 56 positions located within 7.5Å of TMR (see all metrics in **Fig S4** and **S5**). Given that the variability in the model’s performance metrics across the five folds was also low for the positions within 7.5Å (see standard deviations at bottom of **Fig 2C**), we selected this subset for subsequent analyses. Finally, to identify the specific mutations that impact resistance, we used a GB Regressor algorithm with the combined set of 570 BRIGHTE and 208 Single-Env datasets (see normalization of the IC_50_ values and performance in **Fig S6**). The Shapley Additive Explanations (SHAP) value was used to quantify the contribution of mutations to the model’s predictive capacity (44). As shown in **Fig 2D**, performance of the GB Regressor model was high. Importantly, it provided a list of mutations, each with an estimated effect on resistance (**Fig S7**). The 14 mutations with mean absolute SHAP values greater than 0.01 (**Data File S4**) were defined as suspected of impacting resistance, and were further tested *in vitro* as described below.

Machine learning algorithms identify mutations that are frequently sampled in the datasets. To identify mutations that are less frequently sampled, we used a probabilistic modeling approach that only considered mutations appearing in less than 5% of samples (see **Methods Section**). The model applies for each mutation the IC_50_ values across the different samples that contain it, and calculates the relative likelihood of that mutation to contribute to resistance (see **Methods Section** and **Fig S8A**). A total of 17 unique mutations were identified by this approach (see **Fig S8B** and **Data File S5**). In addition, to account for mutations that are not represented in our datasets, we used the structure of the TMR-bound Env (5U7O) to identify Env positions with side chains that extend into the CD4-binding pocket (**Table S2**). We then examined the frequency of all residues at these positions in HIV-1 clade B viruses, represented by a panel of 2,535 Envs from fostemsavir-untreated individuals (see alignment in **Data File S6**). Variants that appeared in at least 0.5% of this panel were selected. A total of 17 such unique mutations were identified. Finally, we identified 11 unique mutations noted in previous studies to increase resistance to TMR or its structural analogs (**Table S3**) (32, 34, 35, 38, 45, 46). The four approaches yielded a total of 59 unique mutations at 21 Env positions that were suspected of increasing Env resistance to TMR (**Fig 2E**).

### Phenotypic analysis of mutations suspected of increasing HIV-1 resistance to TMR

To determine the effects of the 59 mutations on Env fitness and TMR resistance, we introduced them individually into the Env protein of HIV-1 strain AD8. This well-characterized clade B strain occupies a closed conformation typical of Tier-2-like primary HIV-1 isolates (47, 48). Pseudoviruses that contain the mutant Envs were tested for their infectivity using Cf2Th cells that express CD4 and CCR5, and values were normalized for viral particle count by the reverse transcriptase activity in each sample (49). All 59 mutants were also tested for their resistance to TMR. In addition, for each mutation, we examined the emergence frequency after treatment in the escape group, as well as the proportion of the 65 participants that required a single nucleotide change to acquire the mutation. As shown in **Fig 3A**, most suspected mutations did not alter resistance to TMR, while several increased it considerably (e.g., L116P). Interestingly, some mutations were significantly enriched in the escape group, most notably 426L and 375N, which appeared in 74% and 38% of these individuals, respectively (**Fig 3B**). To better understand the basis for the differential emergence frequencies of mutations after treatment, and based on the observed variability in the IC_50_ values, we focused on the subgroup of 18 mutations that increased the IC_50_ by 3.5-fold or more (see shaded region in **Fig 3A**). We designate this subgroup resistance-enhancing mutations (**REMs**). For some REMs (e.g., 375H, 375M and 375Y), their low frequency in the escape group could be explained by the number of nucleotide changes required (see color of data points in **Fig 3A**). However, several poorly sampled REMs (e.g., 375I) only required a single nucleotide substitution. REM emergence frequencies were not associated with their effects on resistance to TMR (**Fig 3C**). By contrast, a threshold effect was observed for the relationship between fitness and emergence rate, whereby low-fitness REMs (e.g., 434K, 255M and 204D) were poorly sampled whereas REMs with favorable fitness profiles and nucleotide substitution requirements showed variable frequencies of emergence.

**Figure 3.**
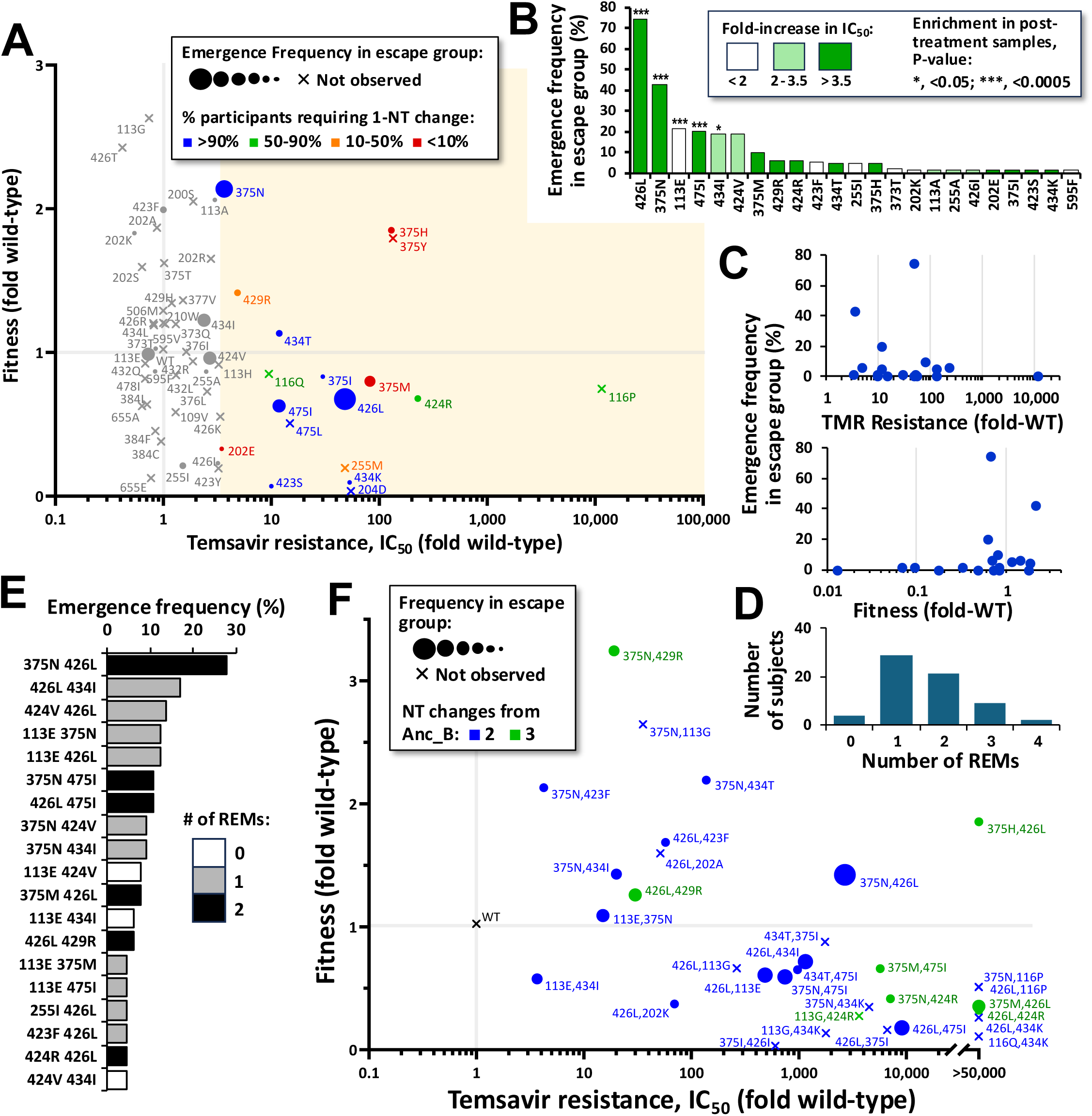
Mutations that increase resistance to TMR are not represented equally in individuals that develop resistance. **(A)** The 59 mutations suspected of increasing resistance to TMR were introduced in the AD8 Env and tested in a pseudovirus system for their fitness (infectivity normalized for virus particle count) and resistance to TMR. Datapoint size corresponds with emergence frequency in the escape group. Color corresponds with the percent of BRIGHTE subjects that required one nucleotide (NT) change to acquire the mutation. **(B)** Emergence frequency of the mutations in the escape group. Significance of mutation enrichment in the post-treatment samples, as determined in a permutation test, is indicated. **(C)** Relationship between frequency of the 18 mutations that increase resistance by more than 3.5-fold (designated REMs) and their effects on TMR resistance or relative fitness. **(D)** Number of REMs that emerged in each of the 65 escape group subjects. **(E)** Emergence frequency of mutation combinations in the escape group. **(F)** Fitness versus TMR resistance of two-mutation combinations. Datapoint size describes their frequency in the escape group, and color describes the number of NT changes required to acquire the mutation from the clade B ancestral form.

In most individuals, REMs appeared at more than one Env position (**Fig 3D**). We thus examined the fitness-resistance profiles of the two-site mutation combinations that appeared frequently after treatment, as well as combinations that appeared less frequently or not at all (**Fig 3E** and **3F**). The most frequent combination in the escape group, 375N/426L, which requires one nucleotide substitution in each codon, exhibited a favorable fitness-resistance profile relative to other two-substitution combinations. Nevertheless, it did not exhibit a unique synergistic advantage in fitness or TMR resistance that could explain the observed high prevalence of this combination or of the individual changes (**Fig S9**).

In the above tests, we compared the emergence frequencies of REMs with their fitness and TMR resistance levels measured using a single HIV-1 isolate (strain AD8). Nevertheless, it could be argued that some poorly sampled REMs, such as 116P, may only increase Env resistance in the context of a limited number of strains (e.g., AD8). Conversely, the 113E and 434I mutations, which appeared frequently after treatment but did not increase AD8 Env resistance significantly, could exhibit limited effects in AD8 relative to other strains. To address this concern, we introduced the 116P, 113E and 434I mutations into the Envs of two clade B transmitted/founder strains, 700010040.C9.4520 and WEAUd15.410.5017 (48). For the 113E mutation, which emerged frequently in BRIGHTE, we also tested the Envs of strains QH_692 and JRFL. For all three REMs, the effects on fitness and resistance were qualitatively similar to those observed for AD8 (**Fig S10**). While these tests included only 2-4 variants, they suggested that the effects of the mutations observed in the Env of strain AD8 can be generalized to other clade B isolates.

In summary, we identified 18 mutations that increase Env resistance to TMR. Some REMs emerged after treatment at significantly higher frequencies than others. This preference could not be fully explained by the number of nucleotide changes required or by the level of resistance to TMR they impart, and only partially by their effects on Env fitness.

### The emergence frequency of REMs in the escape group corresponds with their spontaneous emergence rate in the population

The 426L mutation, which appeared in 74% of subjects in the escape group, increased TMR resistance by 48-fold, while four other suspected REMs at this position (Ile, Thr, Lys and Arg) did not increase resistance significantly (**Fig 3A**). To assess the comprehensiveness of our screening approach, we examined if additional variants at this position may also increase resistance to TMR. To this end, we performed saturation mutagenesis to test the effects of all possible mutations at position 426 on Env fitness and resistance to TMR (50). Replication-competent libraries of HIV-1_AD8_ that contain a degenerate codon at this position were used to infect the T cell line A3R5.7 in the absence or presence of TMR (250 nM). The frequency of each form in the infected cells was compared with the frequency in the virus library used for infection (see protocol in **Methods Section**). While several residues at this position showed fitness levels similar to the wild-type Met, remarkably, only Leu was able to infect the cells in the presence of this concentration of TMR (**Fig 4A**).

**Figure 4.**
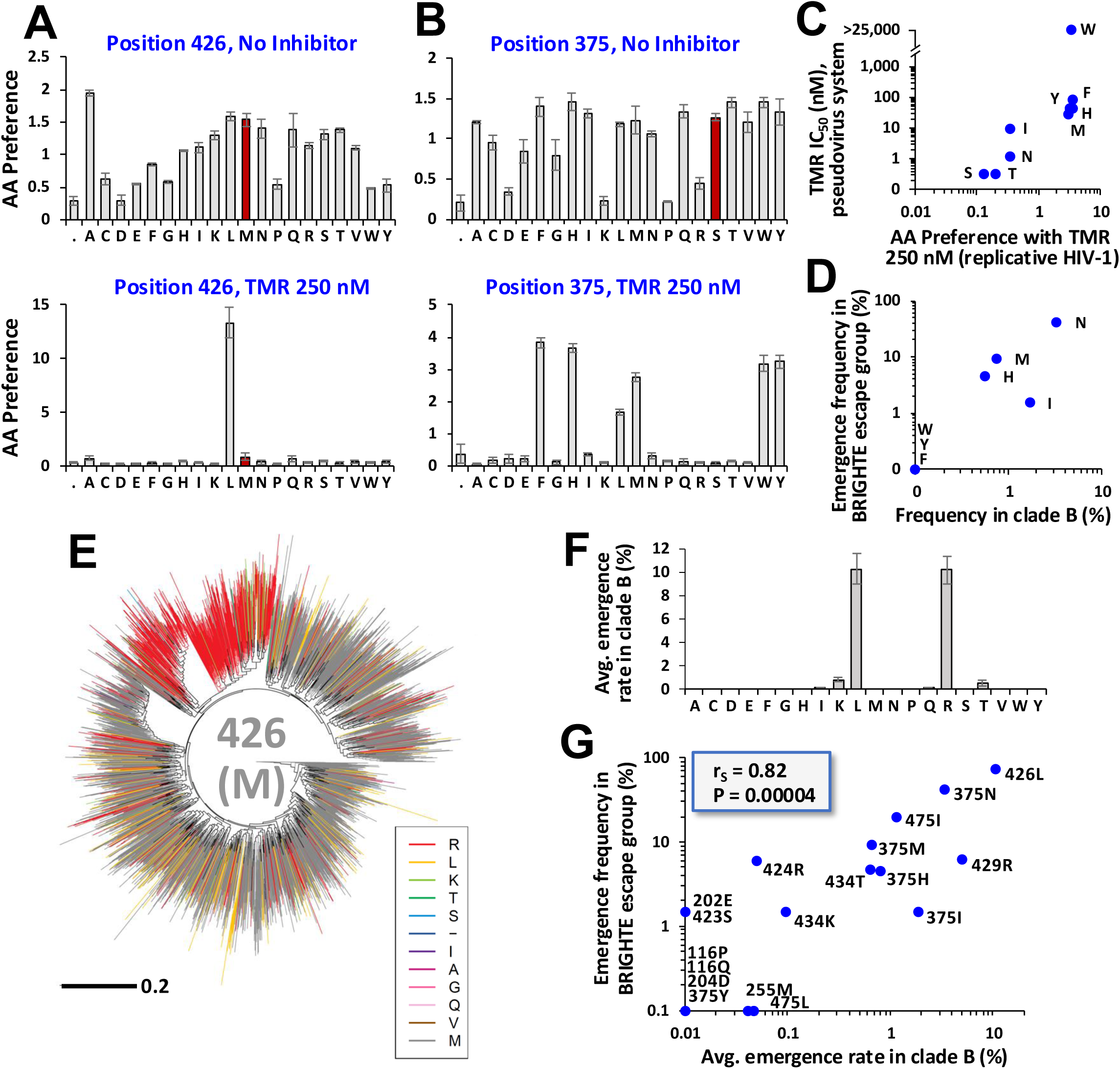
The frequency of REM emergence in the escape group corresponds with their spontaneous emergence frequency in the population. **(A,B)** Saturation mutagenesis to determine the effects of all amino acid changes at positions 426 and 375 on HIV-1 fitness (“No inhibitor”) and resistance to TMR (250 nM). The wild-type form is shown in maroon. **(C)** Relationship between the preference for amino acids at position 375 in the presence of TMR and their IC_50_ values measured using the pseudovirus system. **(D)** Relationship between the emergence frequency of amino acids at position 375 in the escape group and their frequency in a panel of 2,535 clade B Envs from Fostemsavir-untreated individuals. **(E,F)** Example of the emergence rate of mutations at position 426 in HIV-1 clade B. The tree was constructed using amino acid sequences of the 2,535 Envs. Branches are colored by the amino acid in each taxon at position 426. The tree was partitioned into subgroups, which were excluded if the dominant form at that position differed from the clade B consensus. For all remaining subgroups, the number of new substitution events at position 426 to each amino acid was calculated, and these values were averaged. Error bars, standard errors of the mean. **(G)** Relationship between emergence frequency of the 18 REMs in clade B and their emergence frequency in the BRIGHTE escape group.

We also tested the fitness and TMR resistance profiles of all residues at position 375 (**Fig 4B**). REMs 375H, 375I, 375M, 375N and 375Y, which showed considerable increases in resistance using the pseudovirus system (**Fig 3A**), also showed high resistance in these experiments (see correlation in **Fig 4C**). Interestingly, in addition, the 375W and 375F variants showed high resistance to TMR. These forms were not identified by our four approaches because they did not appear in the sequence-IC_50_ datasets or in the panel of 2,535 clade B isolates from fostemsavir-untreated individuals. Indeed, the emergence frequency of mutations at position 375 in BRIGHTE participants correlated well with their proportion in the untreated clade B population (**Fig 4D**). To quantify the propensity for spontaneous emergence of the 18 REMs in fostemsavir-untreated individuals, we applied the panel of 2,535 clade B Envs. The sequences were used to construct a phylogenetic tree that was partitioned into subgroups. Each position was tested separately, by assigning the taxa their amino acid occupancy at that position, and subgroups dominated by the non-ancestral form at that position were excluded (see example of position 426 in **Fig 4E**). The average number of independent emergence events of each REM and the variability across the subgroups was then calculated (see **Fig 4F**). A strong correlation was observed between the average rate of spontaneous REM emergence in clade B and their emergence in the escape group (see **Fig 4G** and **Fig S11**). These findings suggested that the same selection pressures exerted in fostemsavir-untreated individuals may determine the likelihood of the REMs to emerge on treatment.

### Immune pressures restrict the escape paths of HIV-1 from TMR

Selection pressures on replicative fitness and inhibitor resistance are primary factors that guide the evolution of virus proteins in treated individuals. Interestingly, the relationships shown in **Fig 3C** only suggested a threshold effect for fitness and none for the level of resistance imparted by the REMs. We thus sought to identify additional factors that may contribute to the observed preference for some REMs to emerge (on treatment and in the clade B population). In most HIV-infected individuals, antibodies are elicited against conserved epitopes that overlap the coreceptor-binding site (CoR-BS), CD4-binding site (CD4-BS), and inner domain of gp120 (51, 52). These epitopes are not exposed on the native state of Env in most primary isolates, and are thus designated non-neutralizing. The non-neutralizing antibody response was suggested to reduce the risk of infection in vaccine trials (53, 54) and alter the evolutionary path of HIV during early stages of infection (55-57). In both cases, the attributed mechanism was based on Fc-mediated effector functions. To examine the potential effect of such antibodies on mutation emergence during treatment, we measured the sensitivity of the REMs to plasma samples from two HIV-positive individuals that do not reduce infectivity of Env AD8 at the lowest dilution tested (1:160). As shown in **Fig 5A**, several REMs that did not emerge on treatment were neutralized by the plasma (e.g., 116P and 434K), whereas REMs that were sampled more frequently were resistant.

**Figure 5.**
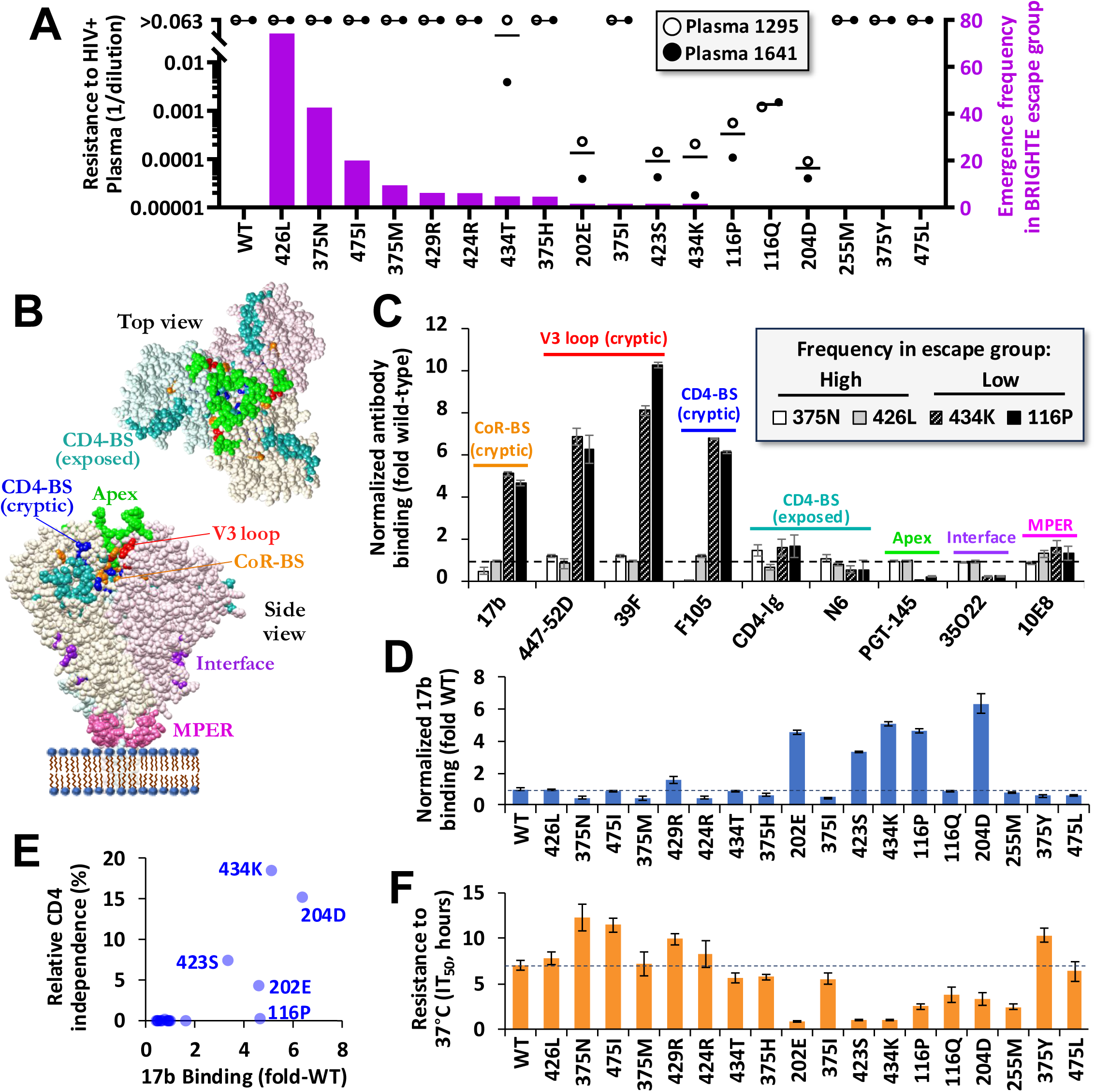
REMs that are poorly sampled in the BRIGHTE escape group exhibit an open conformation of Env that is sensitive to non-neutralizing antibodies. **(A)** Sensitivity of the 18 REMs to plasma from two HIV-infected individuals. Their emergence rates in the escape group are shown as magenta bars. **(B)** Cryo-EM structure of the Env trimer ectodomain in the CD4-unliganded state (PDB IDs 6U59 and 6UJV). Residues associated with binding of the antibodies we tested are colored. **(C)** Envs containing the indicated REMs were expressed on HOS cells, and binding of monoclonal antibodies was measured by cell-based ELISA. Values were normalized for cell surface expression of the Envs using antibody 2G12, and are expressed as a fraction of their binding to the wild-type AD8 Env. **(D)** Binding efficiency of the 18 REMs to antibody 17b that targets a cryptic epitope overlapping the CoR-BS. **(E)** Relationship between binding of the REMs to 17b and their infection of CD4-negative cells, expressed as a percent of their measured infection of CD4-positive cells. **(F)** Resistance of the variants to incubation at 37°C. Values indicate the time until a 50% decrease in infectivity is detected.

To determine the epitope specificity of the antibodies in the plasma that differentially neutralize these mutants, we performed a preliminary experiment using the frequently-sampled (plasma-resistant) 426L and 375N mutants, and the poorly-sampled (plasma-sensitive) 116P and 434K mutants. Plasmids that encode the Envs were used to transfect human osteosarcoma (HOS) cells, which express on their surface fully-cleaved Env trimers in their native closed form (58). Monoclonal antibodies that target different Env epitopes (**Fig 5B**) were added to the cells, and their binding efficiency was detected by cell-based ELISA (59). As shown in **Fig 5C**, the 116P and 434K changes enhanced binding of antibodies against otherwise-cryptic epitopes that overlap the CoR-BS, V3 loop and CD4-BS. In addition, these changes reduced binding of antibody PGT145 that targets a quaternary epitope at the apex of the trimer (60). These features are consistent with a CD4-bound-like conformation of Env (61, 62). We extended the analysis to all 18 REMs and observed that, in addition to 116P and 434K, mutations 423S, 202E, 204D, which appeared infrequently in the escape group and were sensitive to non-neutralizing plasma (**Fig 5A**), also increased binding of the CoR-BS antibody 17b (**Fig 5D**). Given the CD4-bound-like conformation of these REMs, we also tested their ability to infect CD4-negative cells (47). Indeed, a strong relationship was observed between this variable and the level of 17b binding (**Fig 5E** and **Fig S12**). Interestingly, only the 116P mutant showed high binding of 17b but no CD4-independent infection, suggesting two potential structural-functional outcomes for these changes.

We also examined the effects of the 18 REMs on Env stability. The Envs of diverse HIV-1 isolates can exhibit different levels of conformational stability (47, 59) and sensitivities to inactivation at physiological temperature (63, 64). Nevertheless, a relationship between stability of Env variants and their likelihood to appear in HIV-infected individuals has not been established. To this end, we incubated viruses containing the 18 REMs at 37°C and measured the changes in residual infectivity over time. Consistent with previous results for the related ADA strain (64), the half-life of the wild-type AD8 Env was approximately 7 hours (**Fig 5F**). Interestingly, the infrequently sampled REMs that were sensitive to non-neutralizing plasma and enhanced exposure of the CoR-BS also demonstrated low functional stability at 37°C (see correlations in **Fig S13**).

Taken together, these results suggest that REMs 116P, 434K, 204D, 202E and 423S increase sensitivity to non-neutralizing plasma by inducing an open (CD4-bound-like) form of Env that exposes cryptic epitopes targeted by non-neutralizing antibodies. These mutations also reduce Env stability and functional fitness. Their enhanced resistance to TMR can be explained by a change in the CD4-binding pocket to a CD4-bound-like conformation that is not conducive with binding of TMR (note location of the SM^3^ sites in **Fig S14**). However, their resistance is associated with a cost – elimination by the non-neutralizing antibody response.

### Amino acid substitution likelihoods impact the path of resistance *in vivo*

We examined the relationships between the above-described features of the 18 REMs, including their emergence frequencies during treatment and in the clade B population (see values in **Fig 6A** and P-values for the Spearman correlation tests in **Fig 6B**). We divide these features into four groups: **(i) Functional fitness**, measured by the fusion competence of the Env, **(ii) Functional stability**, measured by resistance to inactivation at 37°C, **(iii) Immune fitness**, captured by exposure of the CoR-BS, sensitivity to the non-neutralizing plasma, and indirectly by the requirement for CD4 to infect cells, and **(iv) Therapeutic fitness**, measured by the *in vitro* resistance to TMR. Strong correlations were observed between the variables that capture functional fitness, stability, and immune fitness. However, none of them alone could fully explain the emergence frequency of the 18 REMs in the escape group, other than a modest correlation with Env stability at 37°C. As such, we decided to generate a combined fitness variable that describes the functional fitness, stability and immune resistance of each REM. It was calculated as the product of their relative infectivity, plasma resistance, and stability at 37°C (expressed as a fraction of the values measured for the wild-type Env). As shown in **Fig 6C** and **6D**, the combined fitness of the REMs correlated well with their emergence rates in the BRIGHTE subjects and in clade B. Yet some REMs, such as 375Y, showed high combined fitness but did not emerge on treatment.

**Figure 6.**
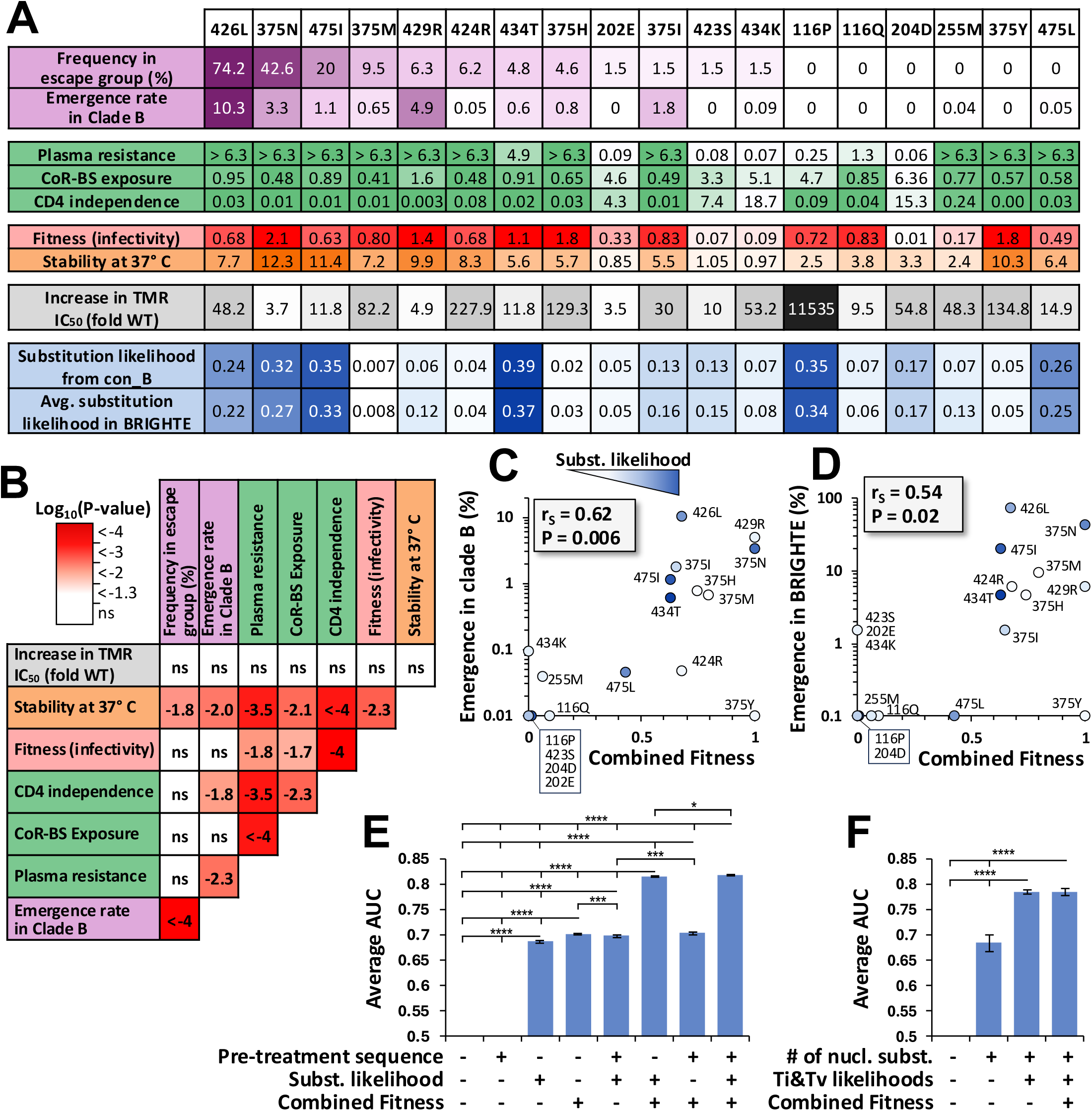
Virus and host factors that guide emergence of the REMs. **(A)** Summary of all features measured for the 18 REMs. Substitution likelihoods from the clade B consensus (con_B) trinucleotide sequence describes the number of nucleotide changes required and the transition/transversion rate for each change. This likelihood was also calculated using the pre-treatment nucleotide sequences of the 65 escape group subjects. **(B)** P-values for a Spearman correlation test that compares the values shown in panel A. ns, not significant. **(C,D)** The combined fitness metric for each REM was calculated as the product of their effects on fitness, resistance to plasma, and stability at 37°C. This value was compared with REM emergence frequencies in clade B or in the escape group. Data points are colored by the substitutions likelihood based on the con_B sequence or the subjects’ pre-treatment sequences, respectively. **(E)** Simulations of REM emergence were performed with different combinations of the indicated variables. The fraction of success events (mutation acquisition) in the 1000 iterations was compared with REM frequency in the 65 subjects (see algorithm in **Fig S15C**). Data for all 18 REMs were compiled to calculate the AUC. Error bars, SDs for 10 simulations. **(F)** Contribution of number and type of nucleotide substitutions to predict emergence of the five REMs at position 375 (H, I, M, N and Y).

Based on these finding, we also examined the likelihood of the mutations to appear, as determined by: **(i)** The pre-treatment nucleotide sequence of the participants, **(ii)** The number of nucleotide changes needed to acquire each REM, and **(iii)** The type of nucleotide changes required, based on the transition and transversion rates determined by Martinez Del Rio et al., (15) (**Fig S15A**). An algorithm that accounts for these variables was generated, in which mutation appearance was modeled as a series of stochastic events with mutation-specific probabilities (see flowchart in **Fig S15B**). As shown in **Fig 6A** (bottom rows), similar REM appearance likelihoods were calculated using the clade B consensus and BRIGHTE pre-treatment sequences as the starting state. We examined if these likelihoods could explain the outliers in **Fig 6C** and **6D** (see colors of datapoints). As expected, several REMs with high fitness but low emergence frequencies (most notably 375Y, 424R and 375I) exhibited low substitution likelihoods.

We extended our analyses to account for the variables associated with both appearance and persistence of each REM (see flow chart in **Fig S15C**). Here, we examined the ability to predict the emergence frequencies of the 18 REMs in the escape group based on their: **(i)** Pre-treatment trinucleotide sequence at these sites, **(ii)** Likelihood for appearance of the REMs (by number and type of the nucleotide substitutions), and **(iii)** The combined fitness metric that incorporates the functional fitness, stability, and immune fitness. As shown in **Fig 6E**, the model that integrated all variables demonstrated good predictive performance (AUC = 0.82) and was robust across multiple cross-validation folds, as indicated by the error bars. The combined fitness variable and the nucleotide substitution likelihoods contributed to the prediction significantly. Consistent with the data in **Fig 6A**, the use of the participants’ pre-treatment sequences did not improve performance relative to the clade B consensus sequence.

Finally, we explored the contribution of the nucleotide substitution likelihoods to the observed profile of mutations in BRIGHTE. To this end, we examined separately the effects of the number of changes required and the probability for each transition or transversion (15). For these tests, we focused on position 375, which contains the largest number of REMs. Interestingly, both the number of changes and the expected rate of each substitution contributed to predictions of REM emergence (**Fig 6F**). For example, position 375 was occupied in most subjects by Ser with the trinucleotide sequence AGT (or less frequently by AGC). Substitutions to Asn (AAT/AAC) or Ile (ATT/ATC) constitute single nucleotide changes (from G to A, or G to T, respectively). However, the G to A transition is more common than the G to T transversion, likely explaining the higher frequency of Asn in the escape group and population relative to Ile despite favorable fitness profiles in both. Indeed, given similar immune and functional fitness levels for the five REMs at position 375 (**Fig 6A**), addition of the selection pressures did not further improve prediction of the on-treatment mutational outcomes (**Fig 6F**). We note that these calculations are not intended to capture the quantitative effects of the variables on appearance or persistence of the REMs. Nevertheless, they clearly demonstrate that the mutational path is explained well by the properties of Env measured *in vitro* and modeled *in silico*.

## DISCUSSION

### The forces that guide the path of HIV-1 escape from a therapeutic in vivo

Through a comprehensive screen, we identified and then tested 59 mutations suspected of increasing HIV-1 resistance to TMR, of which 18 REMs were found to be significantly impactful. Analysis of the utilization of these paths in individuals treated with fostemsavir revealed that some were highly preferred, whereas others, despite favorable fitness and resistance profiles, were sampled infrequently or not at all. Given that the emergence frequency of the REMs in BRIGHTE corresponded well with their spontaneous appearance rates in fostemsavir-untreated individuals, we hypothesized that additional selection pressures may restrict some paths of escape. Indeed, several REMs poorly represented *in vivo* exhibited “open” forms of Env; their mechanism of resistance is based on induction of a CD4-bound-like conformation that is not conducive with binding of TMR. However, the penalty associated with this path is exposure of otherwise-cryptic epitopes that overlap the CoR-BS and V3 loop. Such forms are readily eliminated by non-neutralizing antibodies that are commonly elicited in HIV-infected individuals (51, 52). For other REMs, most notably at position 375, their poor representation in the escape group was explained by the low likelihood of the substitutions to occur, due to the number or type of nucleotide changes required. Thus, we quantitatively capture the *in vivo* factors that impact the two phases of virus escape from a therapeutic: **(i)** *Appearance* of the mutation, as determined by the number and type of nucleotide substitutions, and **(ii)** *Persistence* of the new variant, as determined by the effects on functional fitness, immune fitness, and stability of the protein. Interestingly, a very high level of resistance imparted by some REMs (e.g., 116P) was not able to overcome low levels of functional and immune fitness. The ability to explain the path of resistance by effects of the individual mutations strongly suggests a limited contribution of adaptive changes to escape of the virus. Indeed, REMs with poor functional or immune fitness were rarely observed *in vivo*. Instead, for up to 5 years of follow-up, the mutations with the most favorable profiles emerged. These findings are consistent with the catastrophic effect of the most readily available mechanism of resistance – transition to a CD4-bound like conformation.

### Role of the non-neutralizing antibody response in restricting escape paths from a therapeutic

An unexpected finding of this study is the effect of non-neutralizing antibodies on the range of mutations that can mediate escape. While the neutralizing response against HIV-1 Env is usually swarm-specific and exhibits limited cross-neutralization with other strains, the specificity of the non-neutralizing response is conserved across different individuals (65, 66). Our results suggest that it indeed constitutes a deterministic pressure that guides the evolutionary path of HIV-1, similar to the requirement for functional fitness. Such pressure is consistent with the inability to isolate CD4-independent primary strains from HIV-infected individuals, despite the clear advantage associated with overriding the need for the CD4 receptor (67). The five REMs that were poorly sampled in the escape group (116P, 434K, 202E, 204D and 423S) induced an open form of Env that exposes the CoR-BS, which likely increased their sensitivity to non-neutralizing antibodies against this epitope in the HIV-positive plasma we used. Interestingly, three of these sites (positions 116, 434 and 204) form a clasp-like structure that appears to stabilize Env in the native untriggered state (**Fig S14C**). Similar interactions across the gp120 and gp41 subunits have been noted to maintain trimer stability, allowing Env to retain the high potential energy required to drive the fusion process (68, 69). Weakening of these interactions results in transitions to lower-energy metastable CD4-bound-like forms which, in the absence of a coreceptor, undergo spontaneous and irreversible conformational changes into non-functional states (70). Indeed, here we observe strong relationships between the functional fitness of the variants, their level of “openness”, and their stability at 37°C. Given the large number of such Env-stabilizing interactions, it is plausible that mutations at other sites, outside the CD4-binding pocket, may also increase resistance to TMR. While these mutations may emerge during *in vitro* selection experiments, they appear to be restricted during *in vivo* escape from this therapeutic.

### Effects of Env structural context on availability of escape paths

Env exhibits tremendous diversity in amino acid sequence within and between hosts (17, 18). Divergent profiles of antigenicity and networks of allosteric interactions result in distinct profiles of sensitivity to therapeutics. In addition, the fitness profile of Env positions may vary between strains, including those derived from the same HIV-1 clade (50). As such, a REM could exhibit high fitness in the context of one strain but low fitness in another, restricting viability of that escape path. Context specificity can limit the ability to infer, for example, from our tests with the AD8 Env to other HIV-1 strains. Here, to better understand the high frequency of two mutations that do not increase TMR resistance in our assays (113E and 434I), and the absence of REM 116P that increases resistance significantly, we tested their effects on additional isolates. Qualitatively similar effects were observed on fitness and TMR resistance of these strains, suggesting that, at least for the CD4-binding pocket, the effects measured for Env AD8 are generalizable. This notion was also supported by the observation that the frequency of most REMs in the escape group was explained well by their appearance and persistence likelihoods based on the *in vitro* data measured for AD8 Env. An exception to this rule is REM 475L, which exhibited a relatively high likelihood for appearance and persistence but was poorly represented in the BRIGHTE escape group. Consistent with this finding, 475L only appeared in one of the 2,535 clade B isolates from fostemsavir-untreated individuals. Comprehensive characterization of the effects of this mutation on Env functional and immune fitness in diverse strains, as well as the potential for other selection forces such as the cytotoxic T lymphocyte response (71, 72), may provide greater insight into the basis for the low emergence rate of this mutation.

### Personalization of antiviral treatments based on virus genotype

Several sites located in the CD4-binding pocket show considerable within-clade variation relative to that expected for the critical function of this domain (**Table S2**). Most prominent are positions 426 and 375, which also harbor the two most frequent REMs. Both sites exhibit broad fitness profiles, as measured by saturation mutagenesis (**Fig 4A** and **4B**). For example, the ancestral Met at position 426 is often replaced by Arg and Leu (**Fig 4F**); both are fit, but only Leu is resistant to TMR. Similarly, several variants at position 375 can impart resistance and exhibit high levels of functional and immune fitness (**Fig 6A**); however, they are restricted by a low likelihood for appearance, due to the number (His, Met and Tyr) or type (Ile) of nucleotide changes required. The frequencies of 426L and 375N in HIV-1 clade B increased at early stages of the AIDS pandemic and seem to have reached steady states, likely reflecting the lack of an advantage relative to the ancestral forms (**Fig S16**). Thus, the prevalence of variants that enhance resistance to TMR, and potentially other inhibitors against the CD4-binding pocket (73, 74), are not low but are also not increasing. Indeed, 24 of the 132 subjects that were sampled before and after treatment showed resistance to TMR prior to initiation of therapy, and 65 developed resistance on treatment (**Fig 1F**). The high emergence frequencies of 426L and 375N reflect their favorable immune and functional fitness profiles and suggest that future inhibitors targeting the CD4-binding pocket should be designed to maintain efficacy against variation at these positions.

A longstanding goal in the field of infectious disease management is development of tools to personalize antiviral therapeutics by rapidly estimating the properties of the infecting virus swarm before treatment (75-78). The non-negligible frequency of TMR-resistant strains in the population and the high rate of resistance that emerges on treatment suggest a need for such tools. Nevertheless, the diversity and complexity of Env can challenge our ability to predict resistance from sequence data (79). As we show here, HIV-1 sensitivity to TMR can be readily inferred from the amino acid sequence of Env, with AUC values exceeding 0.95. Given the limited effects of Env context on these predictions, our algorithms can be applied to strains from diverse HIV-1 clades. An example of the relationship between the predicted and measured IC_50_ values using the GB regressor model is shown in **Fig 2D**. For predicting resistance above 50 nM TMR, the false positive rate was 3% and the false negative rate was 7.1%. Notably, the linearity of the relationship between the probabilities for resistance and the measured IC_50_ values allow us also to consider the estimated level of resistance during clinical decision making. Such performance is high relative to previous analyses of epitopes targeted by BNAbs (79-81), which may be attributed to the relatively low complexity of the TMR epitope, or the restricted range of mutations that can impart resistance and persist in the host.

The sequence data derived from the PhenoSense GT assay and applied in this study does not capture the full diversity of plasma virus variants in each participant. Indeed, genotypic profiling assays used clinically are generally limited to detection of mutations with frequencies greater than 20% of all sequences (82, 83). We expect that sequencing of plasma virus at greater depth will provide the accuracy required to predict initial clinical responses as well as short-term emergence of resistant forms from low-frequency circulating variants. Processing of clinical samples by deep sequencing followed by sequence-based estimation of resistance, as described here, can be achieved with limited expense in short time frames relative to phenotypic *in vitro* assays. The approach has the potential to improve both the time-to-treatment initiation and the efficacy of personalized TMR treatments. Furthermore, if resistant forms are not detected in the pre-treatment sequences, the data can be used to quantitatively describe the likelihood for emergence of each resistance mutation, based on the nucleotide substitution requirements and the estimated effects on functional and immune fitness of the virus.

## METHODS

### Processing and analysis of sequence data

All BRIGHTE trial samples used in this study were collected and analyzed by informed consent. Samples were tested for viral loads and CD4 counts. In addition, some samples were analyzed using the Monogram Biosciences PhenoSense GT assay. Nucleotide and amino acid sequences for 580 samples from 371 subjects were available for analysis. TMR resistance values were available for 570 of these samples. Multiple Env positions contained ambiguous nucleotide or amino acid designations, which reflect the presence of more than one sequenced variant in the donor. To align such sequences, we initially demultiplexed the variants by retaining for each position the amino acid or nucleotide found in the clade B ancestral sequence, or, if not present, the first variant listed at that position. We then used a custom Python code to run MAFFT 7.520 for alignment (84). Sequences were then used to determine their clade associations using the Recombinant Identification Program (RIP) tool (85). Sequence variability information was subsequently reintegrated into the samples to allow more than one form at each position, and sequences trimmed to the subset of 856 positions according to HXBc2 numbering (42).

### The Single-Env dataset and transformation of TMR resistance values

We also used a previously published dataset composed of 208 Envs from diverse clades, each associated with an amino acid sequence and TMR resistance value measured *in vitro* (38). Accession numbers (listed in **Data File S3**) were used to download the sequences from the Los Alamos National Lab (LANL) database (86), which were then aligned and processed as above. TMR IC_50_ values for the Single-Env set were measured using an in-house assay (38), whereas the BRIGHTE trial samples were analyzed using the Monogram Biosciences Entry assay. To allow us to combine the datasets, we converted the Single-Env set values to a distribution similar to that of the pre-treatment samples from the BRIGHTE trial (**Fig S6A**). For this purpose, we first examined the distribution type of the BRIGHTE and Single-Env datasets, including Beta, Lognormal, Exponential, Gamma, and Pareto. Based on the sum of squared errors (SSE) metric, both datasets conformed best to a Beta Distribution. We then converted the Single-Env set IC_50_ values to the distribution of the pre-treatment BRIGHTE samples using a probability integral transform approach. In brief, we first mapped the Single-Env set values to their cumulative probabilities, then applied the inverse cumulative distribution function of the BRIGHTE Beta distribution, and finally rescaled to the target support, ensuring the values follow the BRIGHTE distribution while preserving their rank ordering.

### Classification algorithms to estimate Env resistance to TMR by sequence

We evaluated different classification algorithms for their performance to estimate TMR resistance by amino acid sequence. For these evaluations, as well as for hyperparameter tuning, we defined an IC_50_ of 50 nM TMR as the threshold to distinguish between sensitive and resistant samples. As input for the model, we first used the amino acids at the 856 positions of Env according to the HxBC2 numbering system. To account for sequence ambiguity in the BRIGHTE data, we used one-hot encoding to convert each position into binary features representing the absence (0) or presence (1) of each amino acid in the sample. In addition, given that some subjects had multiple samples analyzed (before and after treatment), we used a group-stratified 5-fold cross validation approach, which ensured that all sequences of each subject are assigned to either the training or test folds.

Following preliminary testing of different classification algorithms, we pursued Extreme Gradient Boosting (XGBoost) from the *xgboost* package in Python (43), for its high performance across several key metrics (**Fig S3**). Parameter fine-tuning was performed through grid search in Python to enhance the model’s predictive capacity. The following hyperparameters were tuned: The number of estimators was set to 100, 150, 200, 250 and 300 trees; learning rate (η) was set to 0.1, 0.2 and 0.3; maximum depth of trees was set to 3, 5 and 8; Lambda Regularization was 1, 2, and 3; and the fraction of features and samples used by the algorithm was 1. The AUC was used as the objective function for optimization. Classification metrics were calculated using the *metrics* module from the *sklearn* library (87).

### Gradient Boosting Regressor to identify Env features that impact resistance to TMR

To identify the sequence features that contribute to TMR resistance, we used the Gradient Boosting Regressor (GBR) algorithm. The algorithm was trained on the combined dataset of 570 BRIGHTE and 208 Single-Env samples. Amino acid sequences were used as the input feature set, and the log_10_-transformed TMR IC_50_ values as the response variable. To prepare the sequences, we first excluded Env positions with a minimal distance greater than 7.5 Å from any TMR atom on the TMR-liganded structure of Env (PDB ID 5U7O), resulting in 56 remaining positions (**Table S1**). Next, we applied one-hot encoding to convert the amino acid occupancy at each position into binary features. Of the 243 remaining features, nine exhibited no variation and were excluded. To further reduce dimensionality and mitigate multicollinearity, we dropped the least frequent feature at each position that meets two criteria: **(i)** The position contains at least two unique AA variants in the dataset, and **(ii)** The mutation is included in the feature set tested by the probabilistic approach (see below). This reduced the feature set to 129.

The GBR model was trained using mean squared error (MSE) as the objective function for optimization with the Friedman improvement score as the criterion for measuring split quality, and a learning rate (𝜂) value of 0.1. A k-fold nested grouped cross-validation strategy was used with 𝑘 = 10 for the outer folds and 𝑘 = 5 for the inner folds. This nested structure separates the parameter optimization process from the model assessment with the inner cross-validation loop for hyperparameter tuning, and the outer loop for model evaluation. The grouping strategy ensured that sequences from the same subject appeared in either the training or test fold, but not both. Hyperparameters of the GBR model were optimized via grid search, including the number of estimators (20, 50, or 100) and the maximum depth of individual regression estimators (3, 5, 10, or full tree). Predicted IC_50_ values that exceeded 5 µM were capped at the maximal allowable value of 5 µM TMR. To estimate the contribution of features to the model’s predictions, we used the Shapley Additive Explanations (SHAP) value (44). SHAP values quantitatively describe the impact of each feature on the model’s performance to predict the outcome, including their magnitude, direction and distribution.

### Probabilistic approach to identify low-prevalence mutations that impact TMR resistance

The occurrence of mutations at a low frequency in the dataset limits the ability of algorithms to model their effect. To estimate their impact, we implemented a separate procedure. Mutations included in this set were defined by three criteria: **(i)** They must appear in at least two unique sequences, **(ii)** They must not appear in more than 5% of all sequences, and **(iii)** The side chain of the position must be located within 7.5 Å of the TMR molecule. This filtering process resulted in 107 mutations. If a mutation appeared in more than one sample of a BRIGHTE participant, the average value of the log_10_(IC_50_) was used.

A prioritization algorithm was used to rank these mutations by their potential impact. For each of the 107 mutations, described as amino acid 𝑎 and position 𝑝, we modeled the log_10_-converted IC_50_ values of samples containing that mutation using a normal distribution 𝑋^𝑎,𝑝^, with average 𝜇 and variance 𝜎^2^. We assume that 𝑋^𝑎,𝑝^ corresponds with the distribution of their effects on the greater population of HIV-1 strains (𝑌^𝑎,𝑝^). To compare between the 107 mutations, we randomly sampled with replacement the distribution 𝑋^𝑎,𝑝^ of each mutation. This value was compared with the randomly sampled value from all other 106 mutations and ranked (highest value received the lowest rank). This process was repeated 10,000 times, and the average ranks for all 107 mutations across all iterations were calculated. **Fig S8B** summarizes the top features identified by this approach.

### Production and testing of pseudoviruses containing the Env mutations

All 59 mutations suspected of increasing Env resistance to TMR were introduced by site-directed mutagenesis into a pSVIII vector that expresses the Env of strain AD8 under control of the LTR promoter (88). Pseudoviruses that contain the variants were generated by transfection of HEK 293T cells. Briefly, 9.5 × 10^5^ cells were seeded in each 6-well plate well, and transfected the next day with 0.4 μg of the HIV-1 packaging construct pCMVΔP1ΔenvpA, 1.2 μg of the firefly luciferase-expressing construct pHIvec2.luc, 0.4 μg of pSVIII expressing HIV-1 Env, 0.2 μg of pRev expressing HIV-1 Rev, and 4.2 μL of JetPrime reagent (PolyPlus). The medium was replaced the next day, and virus-containing supernatant was collected 24 h later. Samples were cleared of cell debris by centrifugation at 800 × g and filtered through 0.45 μm pore sized membranes.

As a measure of virus particle content, we quantified the reverse transcriptase activity in the samples using a modified version of a previously published protocol (49), where TaqMan chemistry was used in place of SYBR Green. In brief, RT-qPCR reactions (25 μl total volume) were prepared with TaqMan Gene Expression Master Mix (ThermoFisher) and contained 2.5 mU of MS2 Bacteriophage RNA (Roche), 1 μM of MS2 Forward Primer (5’-TCCTGCTCAACTTCCTGTCGAG -3’), 1 μM of MS2 Reverse Primer (5’-CACAGGTCAAACCTCCTAGGAATG -3’), 200 nM Probe (6[FAM] CGAGACGCTACCATGGCTATCGCTGTAG [TAMsp]), 0.5 U RNase Inhibitor (Fermentas, EO0381), and 2 μl of pseudovirus sample. Pseudovirus samples were lysed in 0.125% Triton X-100, 50 mM KCl, 100 mM Tris HCl (pH7.4), 0.4 U/μl RNase Inhibitor, and 20% glycerol. A standard curve was generated using a dilution series of recombinant HIV reverse transcriptase (NIH AIDS Reagent Program, Cat. No. 12583) ranging from 10⁴ to 10¹² pU/μl prepared in the same lysis buffer. Reactions were run in 0.1 ml MicroAmp plates (Applied Biosystems, 4306737) on a Quant Studio 3 Real-Time PCR System under the following cycling conditions: 42 °C for 20 min, 50 °C for 2 min, 95 °C for 10 min, followed by 50 cycles of 95 °C for 15 s and 60 °C for 1 min. Virus infectivity was expressed as the mean luciferase activity measured for each virus stock (in relative light units, see below) divided by the reverse transcriptase activity in that sample.

To measure sensitivity of the variants to TMR (BMS-626529, MedChemExpress) or plasma from HIV-infected individuals, Cf2Th-CD4^+^CCR5^+^ cells were seeded in 96-well opaque white plates at 2 × 10^4^ cells per well and infected the next day. For neutralization assays, pseudovirus samples were pre-incubated with the TMR or plasma for one hour at 37°C. Samples were then added to the target cells and incubated for 3 days in a 37°C 5% CO_2_ incubator. To measure infection, the medium was removed, 35 μL passive lysis buffer (Promega) were added, and samples subjected to three freeze-thaw cycles. To measure luciferase activity, 100 μL of luciferin buffer (15 mM MgSO_4_, 15 mM KPO4 [pH 7.6], 1 mM ATP, and 1 mM dithiothreitol) and 50 μL of 1 mM d-luciferin potassium salt (Syd Labs, MA) were added to each sample. Luminescence was recorded using a Synergy H1 microplate reader (BioTek).

To measure the decay rate of virus infectivity at 37°C, pseudovirus stocks were divided into aliquots (one sample for each time point), and all samples were snap-frozen on dry ice immersed in ethanol and stored at −80°C. At different time points, samples were thawed in a 37°C water bath for 2 min and then further incubated at 37°C for 2-48 hours. All samples were subsequently added to the Cf2Th-CD4+CCR5+ cells and infectivity measured 3 days later by luciferase activity. All tests of infectivity and neutralization were performed in at least three independent experiments that contain three replicates each. Standard errors of the mean were used to quantify the variability in the data.

### Statistical analysis of mutation enrichment in the escape group after treatment

We compared the frequency of all 59 suspected mutations in the samples collected before and during treatment in the escape group. The escape group was defined as participants for whom all pre-treatment samples had IC_50_ values lower than 50 nM TMR, and at least one on-treatment sample with an IC_50_ greater than 50 nM. To determine enrichment of mutations after treatment in these 65 participants, we tested the null hypothesis that the frequency of the mutations in the pre-treatment samples was equal to or greater than their frequency in the post-treatment samples. Briefly, for each subject we merged all amino acids at each position into a single pre-treatment sample or post-treatment sample. We then calculated for each of the 59 suspected mutations the ratio between the mutation frequency in the post-treatment samples relative to the pre-treatment samples. If a participant contained a mutation in both pre- and post-treatment samples, the individual was excluded from the analysis of that mutation. Timepoint identifiers were then permuted 10,000 times and for each iteration the ratio was calculated. To avoid dividing by zero, a small constant value was added to the denominator. The fraction of iterations in which the ratio for the permuted data was the same or larger than the non-permuted data was applied as the P-value for this one-sided test.

### Rate of independent substitution events in HIV-1 clade B

We calculated the emergence frequency of the 18 REMs in HIV-1 clade B. To this end, we downloaded Env amino acid sequences from the LANL database, and aligned them to the HXBc2 strain. All sequences that contain any ambiguity or stop codons were removed, as well as sequences within 0.05 amino acid substitutions per site from any other sequence. The resulting dataset consisted of 2,535 Env sequences. We note that while no information was associated with these sequences to indicate lack of fostemsavir treatment of the hosts, the sample collection dates (99.2% collected before FDA approval of fostemsavir) and the sources of the remaining samples render the likelihood of such a coincidence negligible. Phylogenetic relationships between sequences were calculated using FastTree (89) on the Galaxy platform (90), and the tree was divided into sublineages comprising 80 to 150 sequences each using the Depth-First Search algorithm (91). All sublineages were processed using HyPhy SLAC (92) on Galaxy to infer the most likely ancestral sequence. A custom python code was then used to calculate the rate of independent mutation events at each Env position using the results of SLAC as an input. In brief, each subgroup tree was recursively searched to determine all substitution events relative to the prior inferred node. A subgroup was ignored if the majority variant differed from the clade consensus sequence. The rate of independent emergence events of for each amino acid was calculated as the ratio between the number of emergence events and the total number of nodes. Daughter nodes of inferred substitution events were excluded from this count unless a new substitution event occurred. The code is available through our GitHub repository at https://github.com/haimlab/Independent_mutation_Tree.

### Cell-based ELISA to measure binding of antibodies to cell-surface Env variants

To measure effects of the mutations on Env antigenicity, we expressed the different variants on the surface of human osteosarcoma (HOS) cells and measured binding of monoclonal antibodies by cell-based ELISA, as we described (7, 59, 93). Briefly, HOS cells were seeded in 96-well plates (1.4×10^4^ cells per well) and transfected the next day with plasmids that express Env, Tat, and Rev, using 60, 11, and 6 ng of each plasmid per well, respectively, and 0.12 μL per well of JetPrime reagent. Background antibody binding was quantified using cells transfected with a pSVIII construct containing a premature stop codon at Env position 46. Three days later, cells were washed in blocking buffer (BB) composed of a tris-saline (TS) buffer (140 mM NaCl, 1.8 mM CaCl_2_, 1 mM MgCl_2_, and 25 mM Tris [pH 7.5]) supplemented with 3% bovine serum albumin and 1.1% skim milk. Cells were then incubated for 45 min at room temperature in BB that contains the antibodies at the following concentrations: 17b and F105 at 5 μg/mL; 39F and 10E8 at 2 μg/mL; CD4-Ig, N6, 447-52D, 35O22, PGT145 and 2G12 at 1 μg/mL. Cells were then washed 6 times with BB and incubated with a horseradish peroxidase-conjugated goat anti-human IgG for 1 h at room temperature. Cells were subsequently washed six times with BB and six times with TS buffer. Antibody binding was measured by chemiluminescence using 35 μL per well of a 1:1 mix of SuperSignal West Pico chemiluminescent peroxide and luminol enhancer solutions (Thermo Scientific) supplemented with 150 mM NaCl on a Synergy H1 microplate reader. To correct antibody binding values to the expression level of each Env, we also measured in each experiment their recognition by antibody 2G12 that targets an exposed epitope on the high-mannose patch of gp120 (94). The antibody-to-2G12 binding ratio was calculated for each Env variant, and the value expressed as a fraction of this ratio measured for the wild-type AD8 Env. Binding assays were conducted at least three times in independent experiments, each with three replicate samples. Standard error values were used to describe the variability in the data.

### Saturation mutagenesis to determine effects on Env fitness and resistance to TMR

To introduce a degenerate codon that encodes for all 20 amino acids (and one stop codon), we used primers that contain the trinucleotide sequence NNK at position 375 or 426 (see schematic of approach and all primers in **Fig S17**). N represents any nucleotide, and K represents G or T. The NNK-containing primers were used to amplify an Env segment using as template the proviral vector pNL4-3 that encodes for the Env of strain AD8 with PrimeSTAR Max DNA Polymerase (Takara). This fragment was then ligated to a second fragment by overlapping PCR to generate a combined fragment that spans the entire *env* gene. This product was then cloned into the env-deleted pNLAD8 vector (GenBank ID PV345784) using In-Fusion assembly (Takara), and the product transformed into Zymo Mix&Go DH5α chemically competent cells. At least 350 colonies for each library were collected and pooled, and plasmids were purified using ZR Plasmid Miniprep-Classic. Two micrograms of the provirus library were then used to transfect 1x10^6^ HEK 293T cells cultured in 6-well-plate-wells using JetPrime reagent. The medium was changed after 4 hours, and 48-hours after transfection the virus was harvested, passed through 0.45 μm filters and treated with 100 U/ml DNase-1 (Roche) for 30 min at 37°C. Samples were the divided into aliquots, snap-frozen using dry ice immersed in ethanol, and stored at -80°C until use. Virus titers were determined by plaque assay after a 96-hour infection of TZMbl-GFP cells (BEI Resources, HRP-20041, contributed by David G. Russell and David W. Gludish) in DMEM/FCS supplemented with 20 μg/ml DEAE dextran.

To eliminate mixed-allele virions from the samples, the above virus libraries were used to infect 2x10^6^ A3R5.7 acute lymphoblastic leukemia T cells in 2 mL at an MOI of 0.003 in RPMI/FCS supplemented with 20 μg/ml DEAE dextran. Cells were resuspended in 4 mL fresh culture medium 24 hours after infection, and 2 days later the virus was harvested and filtered as above. Titers were determined using TZM-bl-GFP cells and the samples used to infect a culture of 1x10^6^ A3R5.7 cells in RPMI/FCS supplemented with 40 μg/ml DEAE dextran at an MOI of 0.05. These infections were performed in the absence or presence of 250 nM TMR. At 16 hours post-infection, cells were pelleted and non-integrated viral DNA was purified using QIAprep Spin Miniprep kit (Qiagen). A 900-bp region encompassing position 375 and 426 was then amplified using PrimeSTAR Max DNA Polymerase. For each biological replicate, an initial 30-cycle amplification was performed in triplicate, the samples pooled, and gel-purified using Zymoclean Gel DNA Recovery Kit (Zymo). If necessary, an additional 8-12 rounds of PCR were performed to obtain sufficient sample for sequencing, which was purified using the QIAquick PCR Purification Kit (Qiagen). To sequence the input virus used for the second round of A3R5.7 cell infection, viral RNA was purified using Quick RNA Viral Kit (Zymo), reverse transcribed using SuperScript IV (Invitrogen), and PCR amplified using the same primers used with the purified product from the infected cells. Finally, all samples were sequenced by Oxford Nanopore Technology (PlasmidSaurus), which provided an average count of 5,000 reads per sample.

Sequence data (in fastq format) were used to calculate amino acid preferences at positions 375 and 426 in the absence and presence of Temsavir based on the method described by Haddox and colleagues (50). This approach calculates the relative frequency of each amino acid in viral DNA isolated from cells infected by the virus library relative to their frequencies in the virus library used for infection. To determine if calculations should be corrected for the error rate in the sequencing reactions, we first examined the incorrect variant calls at the two positions by sequencing three samples infected by the wild-type virus. The average frequency of minority variants across the six samples was 0.13% (standard deviation 0.12%). As such, we performed subsequent calculations in an error-agnostic manner.

To calculate the enrichment ratio (𝜙) for each amino acid 𝑎 at position 𝑝, we used:

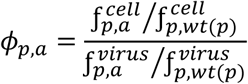

where 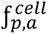 and 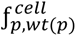 are the frequencies in the cell lysate of amino acid 𝑎 or the wild-type (𝑤𝑡) amino acid for position 𝑝, and 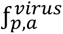and 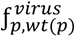 are the frequencies of these forms in the input virus sample used for infection. Finally, we calculated the amino acid preference (𝜋) for each amino acid as:

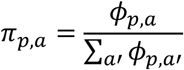

where the 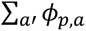 is the sum of enrichments for all amino acids 𝑎′ at position 𝑝.

A complete software package to calculate amino acid preferences is available through our Github repository at https://github.com/haimlab/EZDMS.

### Algorithms to estimate appearance and persistence of REMs

To evaluate the contribution of the different variables to the observed mutational profiles in treated individuals, we designed two algorithms (see flowcharts in **Fig S15B** and **S15C**). In both, mutation appearance and persistence were modeled as stochastic events with mutation-specific probabilities. The first algorithm calculated for each of the 18 REMs individually the probability of appearance based on the number and type of nucleotide changes required from the initial trinucleotide sequence. To test the probability for REM appearance in clade B, the algorithm was initiated with the trinucleotide sequence at the Env position in the clade B consensus sequence. To test the probability for REM appearance in the 65 participants of the escape group, the algorithm was initiated with the trinucleotide sequence in the pre-treatment samples of the 65 escape group participants. One of the three nucleotide sites was selected randomly, and a nucleotide substitution was randomly introduced at that site based on the transition and transversion likelihoods determined by Martinez Del Rio et al., (15) (see values in **Fig S15A**). Up to two consecutive mutations were allowed. If the desired REM was acquired within the two attempts, a success event was recorded. For each REM, the algorithm was repeated 1,000 times for the clade B consensus sequence, and 1,000 times for each escape group participant. The fraction of success events of the 1,000 iterations in each participant was defined as their probability for mutation appearance.

The second algorithm (**Fig S15C**) introduced an additional module to the mutation appearance step, which aimed to capture the ability of each REM to persist in the host. It was based on the *in vitro* measured effects of the REMs on: **(i)** Infectivity, **(ii)** Functional stability at 37°C, and **(iii)** Resistance to non-neutralizing plasma. The three values measured for each REM were expressed as a fraction of the wild-type AD8 Env values, and their product was defined as the Combined Fitness value of each REM. For each success event from the appearance module, persistence was determined by a random process, in which the Combined Fitness value was used as the probability of the REM to persist. If the mutation persisted, a success event was recorded. The fraction of success events of the 1,000 iterations was defined as the probability for REM emergence. Performance was evaluated by compiling the probability values for the 18 REMs in the 65 participants of the escape group, which were compared with their outcomes: the absence (0) or presence (1) of REM emergence in the participant after treatment. Performance was measured using the AUC metric. The code for the above algorithm can be found on our Github repository at https://github.com/haimlab/REM-Emergence-Modeling.

## ACKNOWLEDGEMENTS

This work was supported by National Institutes of Health (NIH) grant R01 AI170205 to HH, and by ViiV Healthcare Investigator Sponsored Study (ID 4104) to HH. The funders played no role in study design, data analysis, *in vitro* data acquisition or analysis, or the decision to publish this work. The authors are grateful to Anthony Roberth Rojas Chávez for assistance with processing of the sequence data.

## SUPPLEMENTAL DATA FILES

**Data File S1.** Amino acid sequence and TMR resistance values for the 570 BRIGHTE trial samples. Sequences were obtained by the Phenosense GT assay. For each sample, the complete amino acid sequence is provided, as well as the sequence at the 856 positions of Env according to the HXBc2 reference strain. TMR IC_50_ values for the samples, as measured using the Phenosense GT assay, are indicated. Clade associations were inferred using the RIP tool.

**Data File S2.** Phylogenetic tree (in Newick format) of 360 pre-treatment samples from BRIGHTE subjects. The tree is rooted to the Env of the HXBc2 reference strain. These data were used to generate the tree in **Fig S2A**.

**Data File S3.** Amino acid sequence and TMR resistance values for the 208 Envs of the Single-Env dataset. Accession numbers reported in Pancera *et al*., (Nat Chem Biol, 2017) were used to download the amino acid sequences of the 208 Envs included in our study. Aligned amino acid sequences are provided for all Env positions and for the 856 positions of Env according to the HXBc2 reference strain. TMR IC_50_ values for the Envs were measured using a TZM-bl neutralization assay.

**Data File S4.** GB regressor model to identify mutations that affect HIV-1 resistance to TMR. Env positions within 7.5Å of the TMR molecule were used as input for the model. SHAP values for all residues at these positions were averaged across all 778 samples used for the prediction. Mean SHAP values as well as the values for all 778 samples are provided.

**Data File S5.** Probabilistic model to detect mutations that impact Env resistance to TMR. Mutations at Env positions within 7.5Å of the TMR molecule were used as input for the model using the 778 samples of the BRIGHTE trial and Single-Env datasets. Only mutations that appear in less than 5% of all samples were analyzed. The average IC_50_ value for all samples that contain each mutation and the variability (measured by the standard deviation) are shown, and were used to rank the mutations according to their likelihood for impacting Env resistance to TMR.

**Data File S6.** Amino acid alignment (in fasta format) of 2,535 Env sequences from HIV-1 clade B.

**Data File S7.** Emergence frequency of amino acid variants at all Env positions in HIV-1 clade B. Values were calculated using the phylogenetic tree constructed from 2,535 amino acid sequences of clade B Envs. They describe the rate of substitution to each amino acid (from the clade ancestral form) at the 856 positions of Env according to HXBc2 numbering. The number of sequences in each subgroup is indicated above each column.

**Data File S8.** Phylogenetic tree (in Newick format) of 2,535 Env sequences from HIV-1 clade B.

