## Supplemental Figures S1-S17 for "Comprehensive Mapping of the Virus and Host Factors that Guide the Paths of HIV-1 Escape from a Therapeutic"

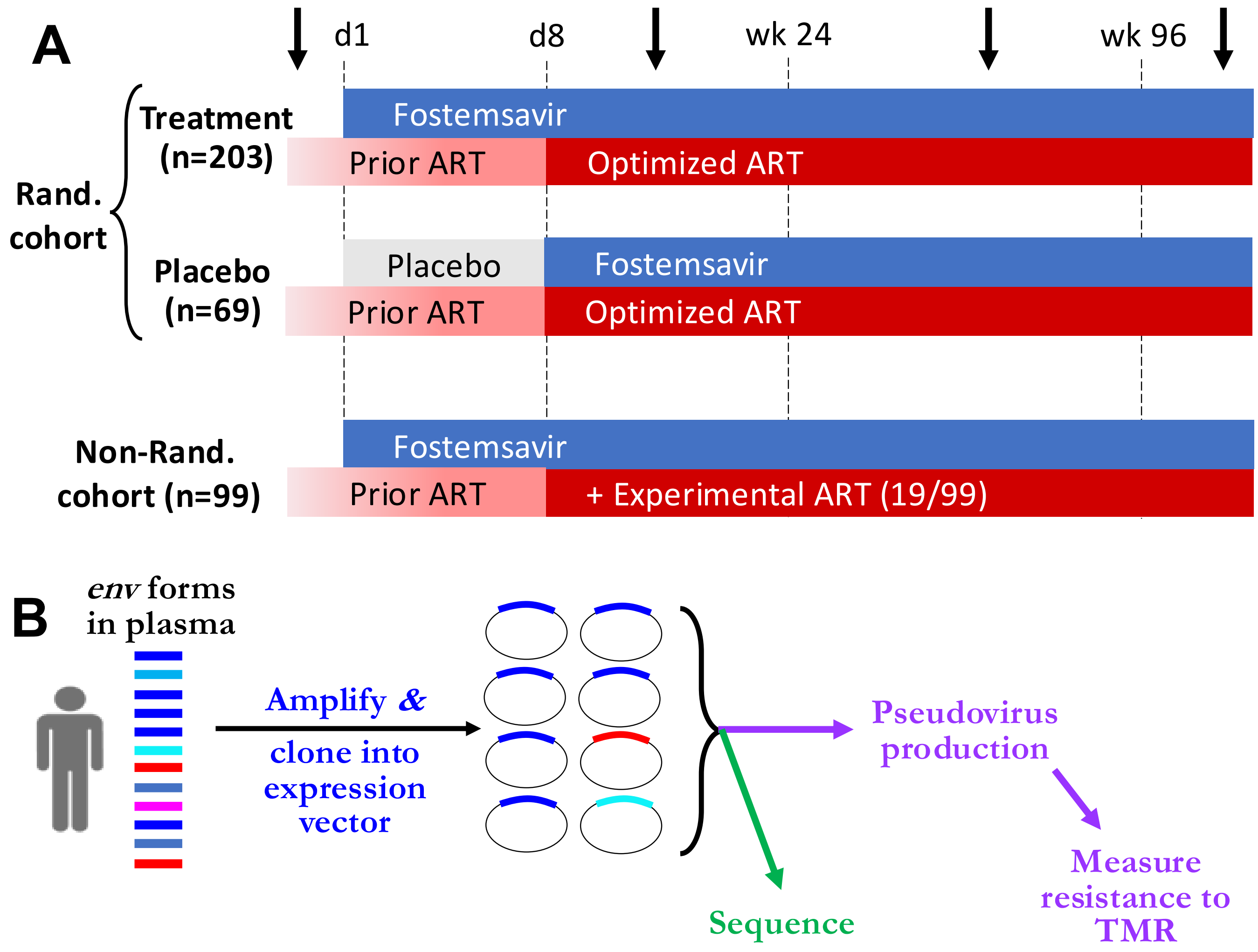


| Subject | Day | 1 | 2 | 3 | 4 | … | 856 | $IC_{50}$ (μM) |
|---|---|---|---|---|---|---|---|---|
| **1** | -46 | M | R | A | K/R | | L | **0.0004** |
| **1** | 239 | M | K | V/A | K | | L | **0.6** |
| **…** | | | | | | | | |
| **360** | -15 | M | T | V | T/K | | L | **0.03** |
| **360** | 64 | M | T | V | T | | L | **>5** |

**Supplemental Figure S1. Design of the BRIGHTE clinical trial. (A)** Prior to trial initiation, phenotyping assays were conducted for plasma samples from all subjects to determine virus sensitivity to different ART classes. Based on these results, subjects for whom a complete ART regimen could not be constructed were assigned to the non-randomized cohort whereas all others were assigned to the randomized cohort. Treatment with Fostemsavir (800 mg bid) or placebo was administered for 8 days, after which this regimen was continued and supplemented by an optimized ART regimen for all subjects of the randomized group. At different time points before and on treatment, blood samples were collected (represented by arrows) and analyzed for viral load, CD4 count, and used for Env phenotyping and genotyping assays. **(B)** The PhenoSense GT assay is based on cloning a library of amplicons containing the *env* gene into an expression plasmid. The plasmid library is then sequenced and used to generate a library of pseudoviruses that is tested *in vitro* for resistance to TMR. Given the in-host diversity of Env, many sites are assigned more than one amino acid, which is addressed by one-hot feature encoding, as described in the Methods Section.

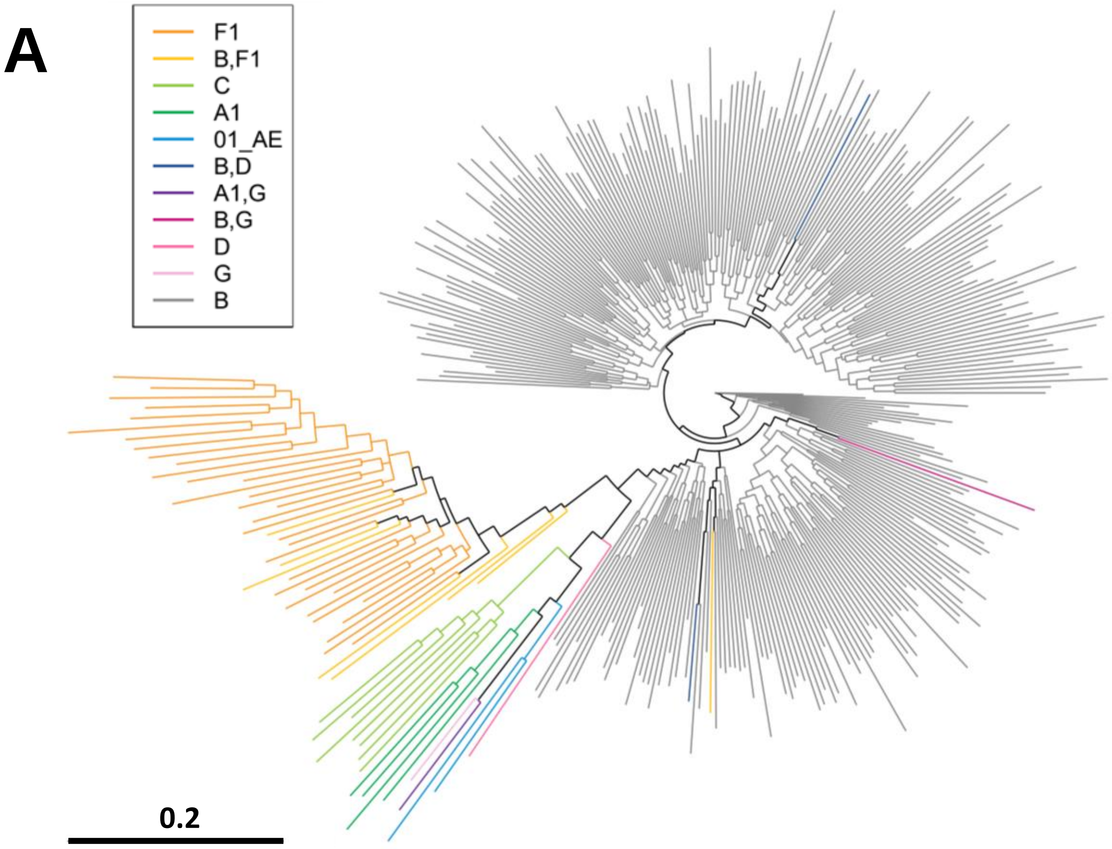


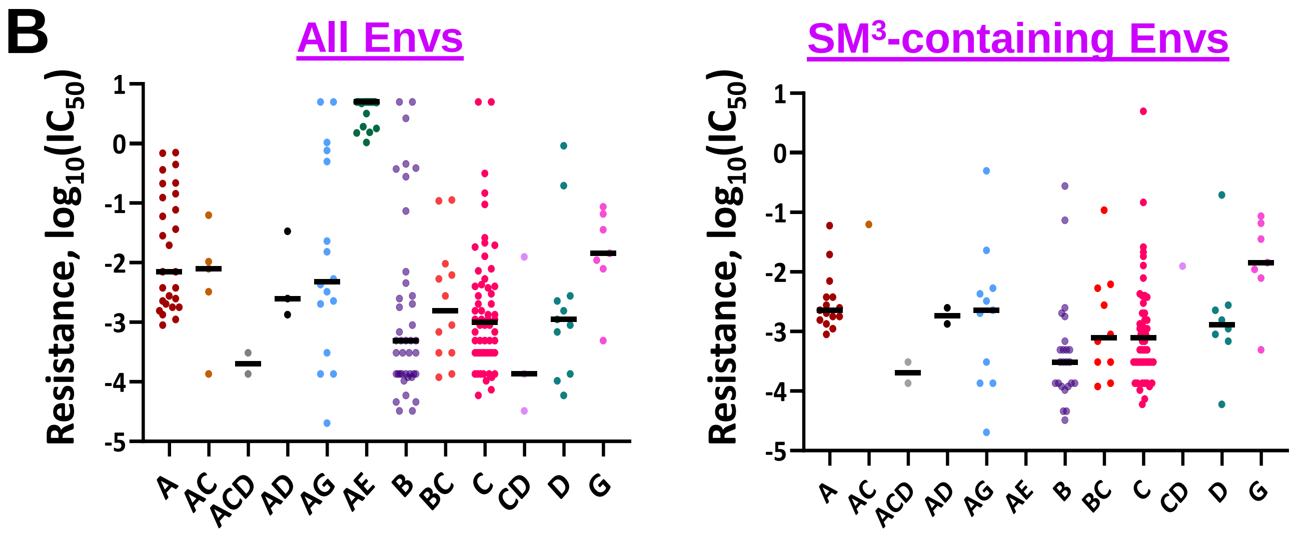


**Supplemental Figure S2. Clade distribution of the BRIGHTE trial dataset and the Single-Env dataset. (A)** Phylogenetic tree based on nucleotide sequences of pre-treatment samples from the 360 BRIGHTE trial subjects. A single sequence was used for each subject. Clade associations were inferred using the RIP tool. **(B)** Resistance values for the 208 Envs of the Single-Env dataset (as reported in Pancera et al., 2017) and for the subset of samples that contains the TMR-sensitive $SM^3$ motif. Samples are grouped by their inferred clade associations as reported by the authors.

| Model | AUC | Accuracy | F1 Score | Precision | Sensitivity | Specificity |
|---|---|---|---|---|---|---|
| **XGBoost** | 0.95 (0.02) | 0.90 (0.03) | 0.82 (0.03) | 0.85 (0.06) | 0.79 (0.05) | 0.94 (0.03) |
| **Gradient Boosting** | 0.94 (0.02) | 0.89 (0.03) | 0.81 (0.04) | 0.86 (0.05) | 0.77 (0.06) | 0.95 (0.02) |
| **Logistic Regression** | 0.92 (0.01) | 0.88 (0.02) | 0.80 (0.02) | 0.83 (0.06) | 0.76 (0.04) | 0.92 (0.04) |
| **AdaBoost** | 0.85 (0.04) | 0.85 (0.03) | 0.79 (0.05) | 0.82 (0.05) | 0.78 (0.09) | 0.88 (0.03) |
| **Random Forest** | 0.79 (0.02) | 0.82 (0.02) | 0.72 (0.03) | 0.75 (0.04) | 0.69 (0.04) | 0.81 (0.01) |
| **SVM** | 0.70 (0.02) | 0.72 (0.03) | 0.70 (0.03) | 0.71 (0.04) | 0.64 (0.02) | 0.73 (0.03) |

**Supplemental Figure S3. Performance of different classification-based algorithms to predict TMR resistance by sequence.** Amino acid sequence data for all 856 positions of Env in the 570 samples from the BRIGHTE trial were used as input to the indicated learners to predict resistance. Resistance was defined as an $IC_{50}$ value greater than 50 nM. Classification metrics are shown. Values in parentheses describe standard deviations for five-fold cross validation. AUC, area under the curve; SVM, Support Vector Machines.

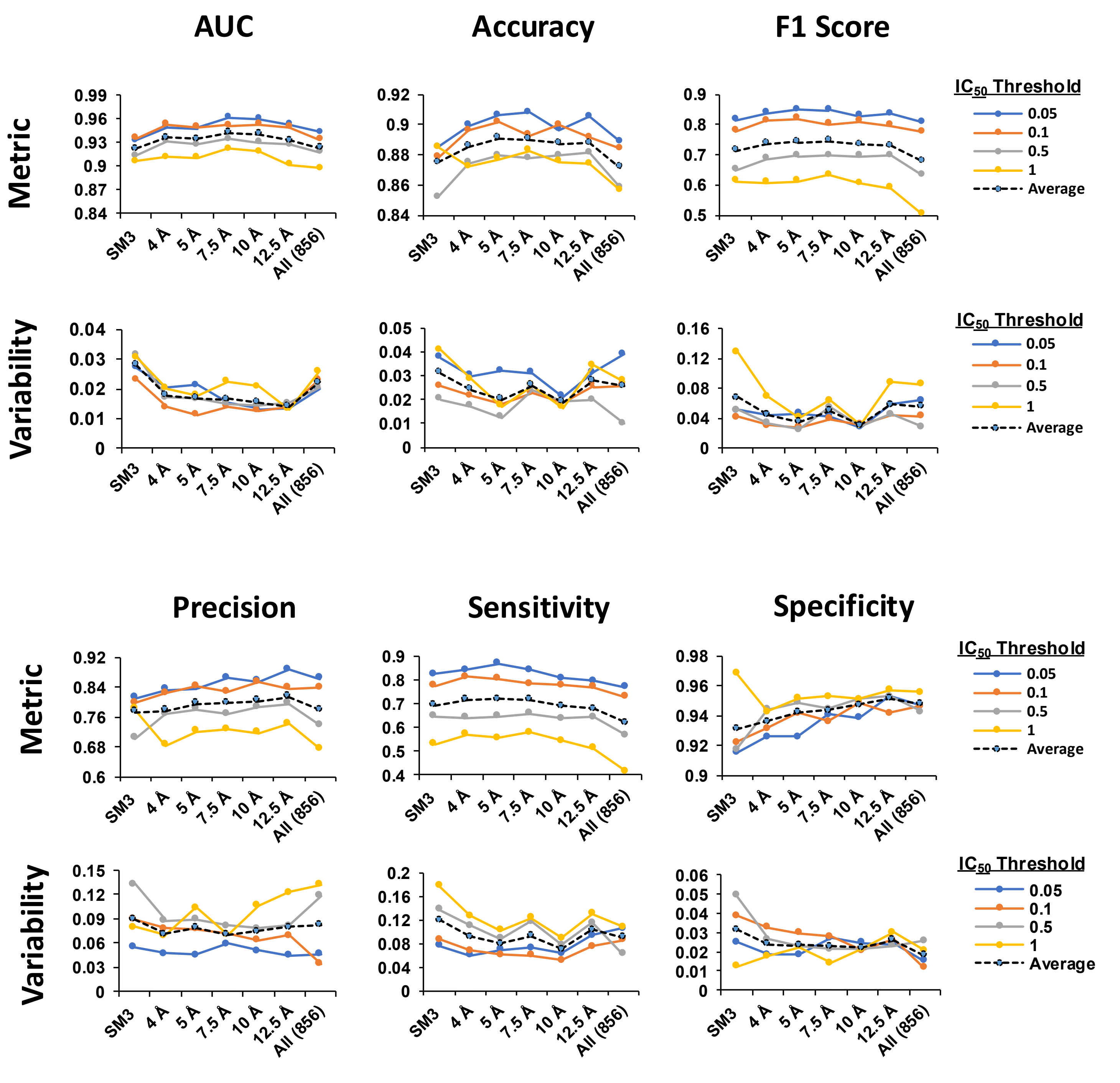


**Supplemental Figure S4. Classification metrics for prediction of resistance for the 570 BRIGHTE samples using the XGBoost algorithm.** As input, we used the amino acid sequences at positions within the indicated distances from the TMR molecule, the four $SM^3$ positions, or all 856 positions of Env. Mean metric values for the five folds are shown in the top graphs and the variability (in standard deviation) in the bottom graphs. Averages values for the different $IC_{50}$ thresholds are shown as black dashed lines.

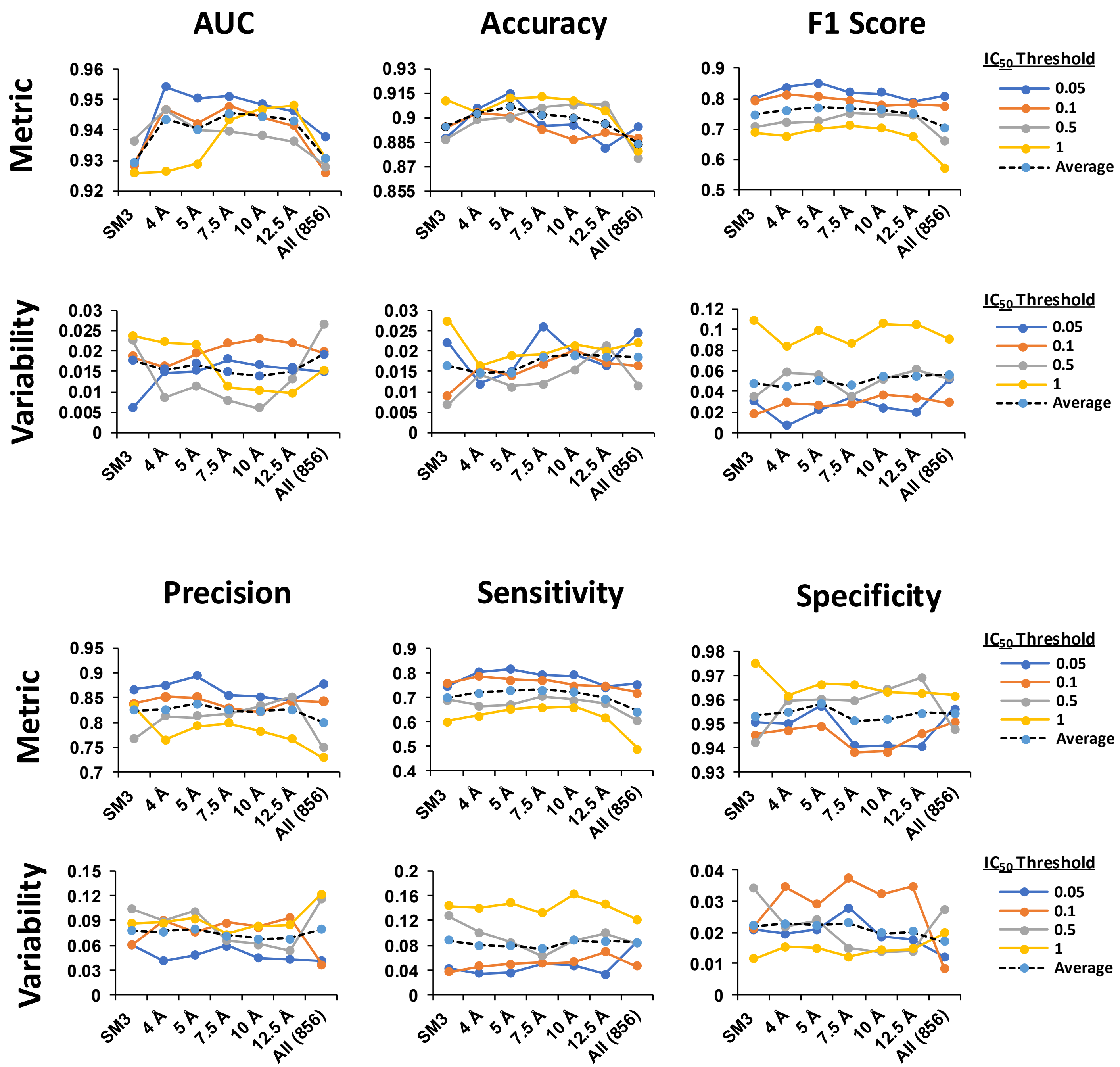


**Supplemental Figure S5. Classification metrics for prediction of resistance using sequence-$IC_{50}$ data from the combined dataset.** Input sequence data included the 570 BRIGHTE samples and the 208 Envs of the single-Env dataset**.** Performance of the XGBoost algorithm was tested as described in **Fig S4**.

A

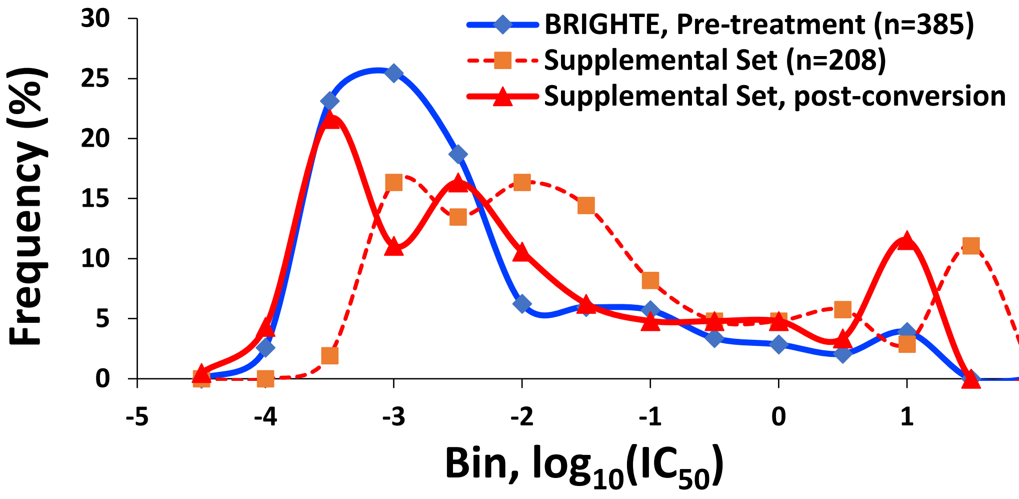


B

| Model | AUC | Accuracy | F1 Score | Precision | Sensitivity | Specificity |
|---|---|---|---|---|---|---|
| XGBoost | 0.94 (0.01) | 0.89 (0.02) | 0.81 (0.05) | 0.88 (0.04) | 0.75 (0.08) | 0.96 (0.01) |
| Gradient Boosting | 0.94 (0.01) | 0.89 (0.31) | 0.80 (0.04) | 0.85 (0.42) | 0.73 (0.07) | 0.94 (0.02) |
| Logistic Regression | 0.87 (0.03) | 0.83 (0.01) | 0.66 (0.05) | 0.85 (0.04) | 0.54 (0.06) | 0.96 (0.01) |
| AdaBoost | 0.89 (0.02) | 0.86 (0.01) | 0.75 (0.03) | 0.81 (0.40) | 0.71 (0.05) | 0.93 (0.01) |
| Random Forest | 0.77 (0.02) | 0.77 (0.02) | 0.54 (0.02) | 0.78 (0.07) | 0.42 (0.10) | 0.80 (0.02) |
| SVM | 0.81 (0.05) | 0.76 (0.02) | 0.68 (0.07) | 0.58 (0.09) | 0.68 (0.06) | 0.85 (0.02) |

C

| Model | AUC | Accuracy | F1 Score | Precision | Sensitivity | Specificity |
|---|---|---|---|---|---|---|
| XGBoost | 0.91 | 0.89 | 0.78 | 0.91 | 0.79 | 0.97 |
| Gradient Boosting | 0.88 | 0.88 | 0.78 | 0.84 | 0.72 | 0.95 |
| Logistic Regression | 0.88 | 0.84 | 0.74 | 0.69 | 0.79 | 0.86 |
| AdaBoost | 0.9 | 0.88 | 0.8 | 0.78 | 0.81 | 0.91 |
| Random Forest | 0.9 | 0.88 | 0.77 | 0.79 | 0.76 | 0.89 |
| SVM | 0.85 | 0.84 | 0.75 | 0.77 | 0.65 | 0.87 |

**Supplemental Figure S6. Combined analysis of the BRIGHTE and Single-Env datasets.** **(A)** TMR resistance values for the BRIGHTE trial samples and the Single-Env dataset were measured using different *in vitro* assays. As such, we normalized the $IC_{50}$ values of the latter dataset to the distribution of these values in the pre-treatment samples from the BRIGHTE trial (see Methods Section). Both datasets conformed to a Beta distribution. **(B)** Performance of the different classification-based algorithms to predict TMR resistance by sequence, using as input for the model the combined 778 samples from the BRIGHTE and Single-Env datasets. $IC_{50}$ values for the latter group were normalized to the distribution of the former, as shown in panel A. Amino acids at the 856 positions of Env according to the HXBc2 numbering system were used as input for the model with a threshold of 50 nM to define resistance. **(C)** Performance of the models to predict resistance of the 208 isolates of the single-Env dataset using a model trained on the 570 samples from the BRIGHTE trial. The amino acid sequence at the 56 positions within 7.5 Å from the TMR molecule on the structure were used as input.

**A**

| Metric | Value |
|---|---|
| MSE | 0.63 ± 0.15[a] |
| MAE | 0.57 ± 0.07 |
| RMSE | 0.26 ± 0.06 |
| RMAE | 0.42 ± 0.05 |

**B**

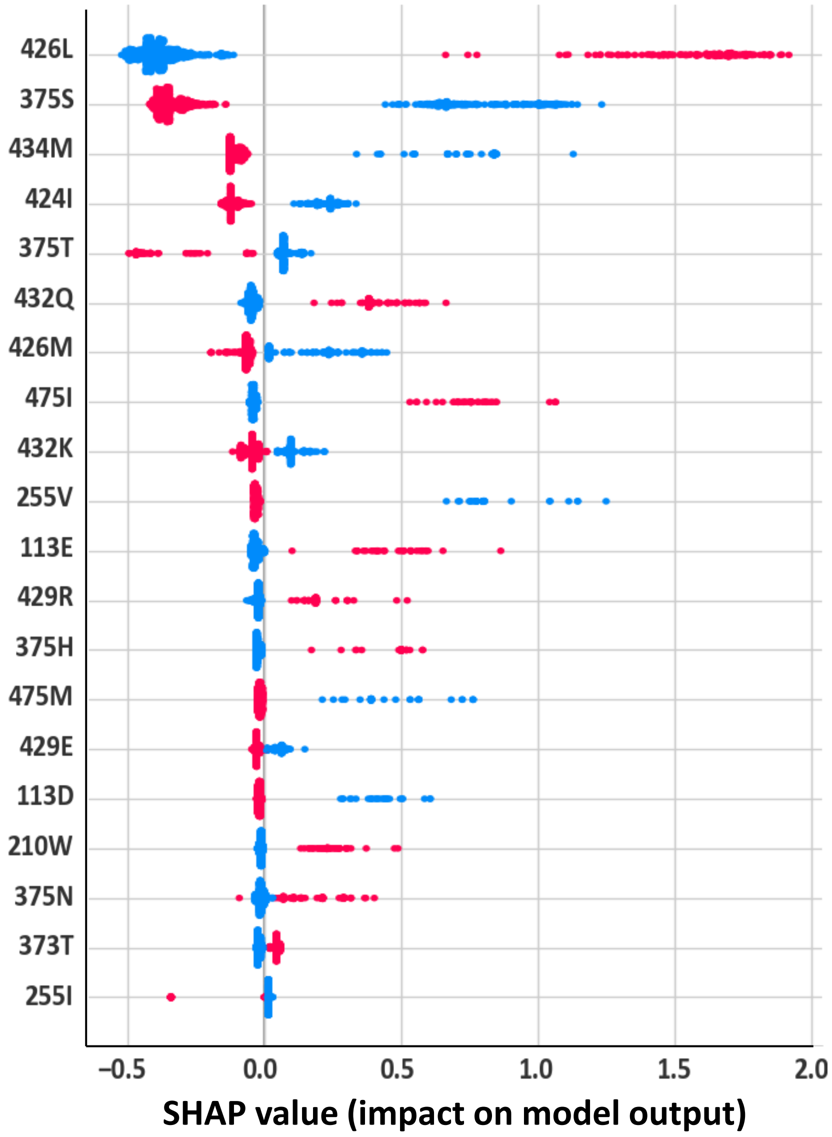


**Supplemental Figure S7. Performance and SHAP values obtained using the GB Regressor model.** The 778 samples of the BRIGHTE and Single-Env datasets were used to predict Env resistance by sequence using the 56 sites located within 7.5Å of the TMR molecule. **(A)** Performance of the model. MSE, Mean squared error; MAE, mean absolute error; RMSE, root mean squared error; RMAE, root mean absolute error. **(B)** Data points describe for each sequence feature the SHAP values calculated for all 778 samples. These values capture the magnitude, direction, and distribution of the impact of each feature on model performance. Red and blue colors describe the presence or absence of the indicated feature in each sequence. Sequence features with the highest SHAP values are shown, and all data are provided in **Data File S4**. Note that many sequence features shown represent the TMR-sensitive amino acid variant.

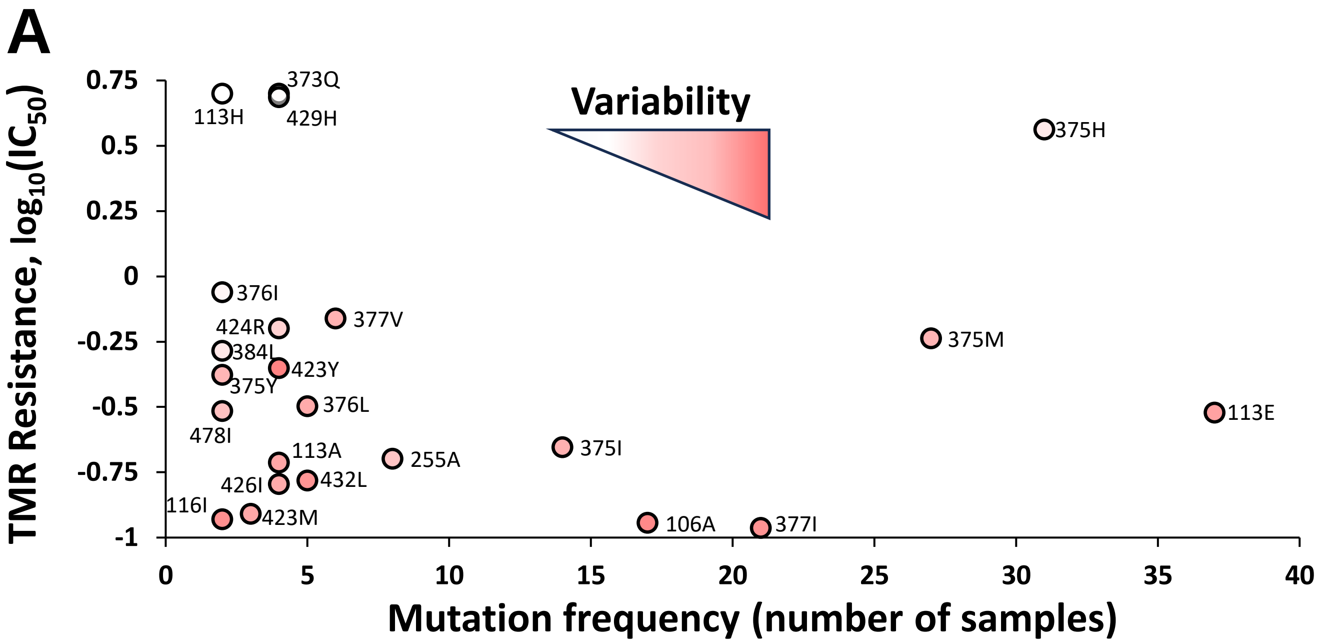


**B**

| # | Feature | Avg. Rank | Distance (Å) | # of unique Seqs ($n$) | $\bar{x}$ | $s$ |
|---|---|---|---|---|---|---|
| 1 | 113H | 2.3 | 1.7 | 2 | 0.70 | 0.00 |
| 2 | 373Q | 2.3 | 6.2 | 4 | 0.70 | 0.00 |
| 3 | 429H | 3.5 | 7.0 | 4 | 0.69 | 0.02 |
| 4 | 375H | 4.9 | 2.5 | 31 | 0.56 | 0.31 |
| 5 | 376I | 9.4 | 3.2 | 2 | -0.06 | 0.12 |
| 6 | 375M | 12.2 | 2.5 | 27 | -0.24 | 1.09 |
| 7 | 424R | 12.2 | 2.9 | 4 | -0.20 | 0.67 |
| 8 | 377V | 12.3 | 5.0 | 6 | -0.16 | 1.14 |
| 9 | 384L | 13.1 | 3.5 | 2 | -0.29 | 0.34 |
| 10 | 113E | 17.1 | 1.7 | 37 | -0.52 | 1.37 |
| 11 | 375Y | 18.0 | 2.5 | 2 | -0.38 | 1.08 |
| 12 | 376L | 18.2 | 3.2 | 5 | -0.50 | 1.23 |
| 13 | 423Y | 18.9 | 6.3 | 4 | -0.35 | 1.82 |
| 14 | 478I | 19.5 | 6.9 | 2 | -0.52 | 0.91 |
| 15 | 375I | 20.0 | 2.5 | 14 | -0.65 | 1.15 |
| 16 | 255A | 20.8 | 2.3 | 8 | -0.70 | 0.87 |
| 17 | 113A | 23.5 | 1.7 | 4 | -0.71 | 1.31 |
| 18 | 426I | 24.6 | 3.6 | 4 | -0.80 | 1.22 |
| 19 | 432L | 25.6 | 2.5 | 5 | -0.78 | 1.56 |

**Supplemental Figure S8. Probabilistic model to detect low-prevalence mutations suspected of increasing resistance to TMR. (A)** For all mutations at sites within 7.5A of TMR, we examined their frequency in the 778 samples from BRIGHTE and Single-Env datasets. These values are plotted against the average TMR resistance of the samples that contain them. Data points are colored by the variability in the $IC_{50}$ values. **(B)** Summary of the rare and potentially impactful features for the 19 mutations with the highest ranks (i.e., highest likelihood for association with resistance). The smallest distance between the closest atoms of the indicated position and the TMR molecule is shown. $\bar{x}$ , the average $\log_{10}$ TMR $IC_{50}$ value among the samples that contain the mutation; $s$, the variability (in standard deviation) among the $\log_{10}$ $IC_{50}$ values of the samples that contain each mutation.

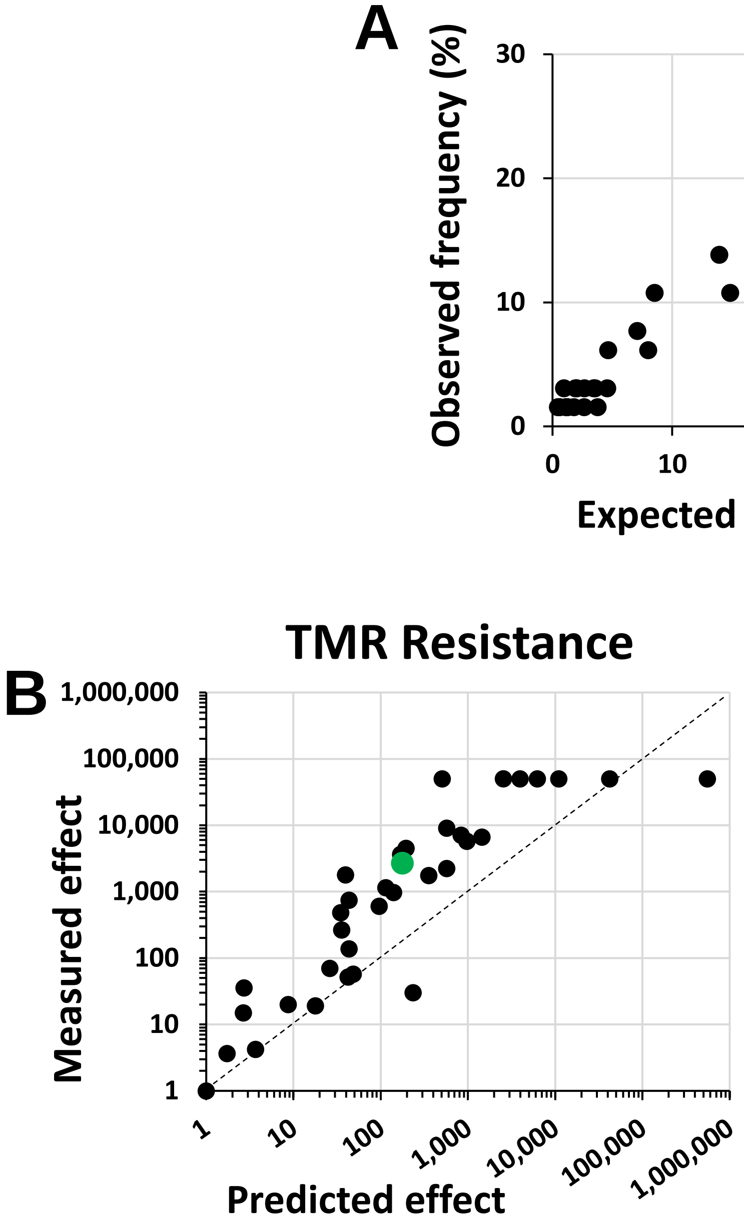


**Supplemental Figure S9. Effects of two-site mutation combinations on Env fitness and resistance to TMR. (A)** The expected frequency represents the product of the individual frequencies of the REMs in the escape group. The observed frequency describes the percent of individuals from the escape group that developed both mutations after treatment. **(B,C)** Synergy between the effects of mutations on Env resistance to TMR and fitness. The measured effect of two-mutation combinations on resistance or fitness (shown in **Fig 3F**) is compared with the predicted effect, which is calculated as the product of the effects of the individual mutations on the respective phenotype. The datapoint for the 375N/426L combination is shown in green. The dashed lines represent the lines of equivalence (x = y).

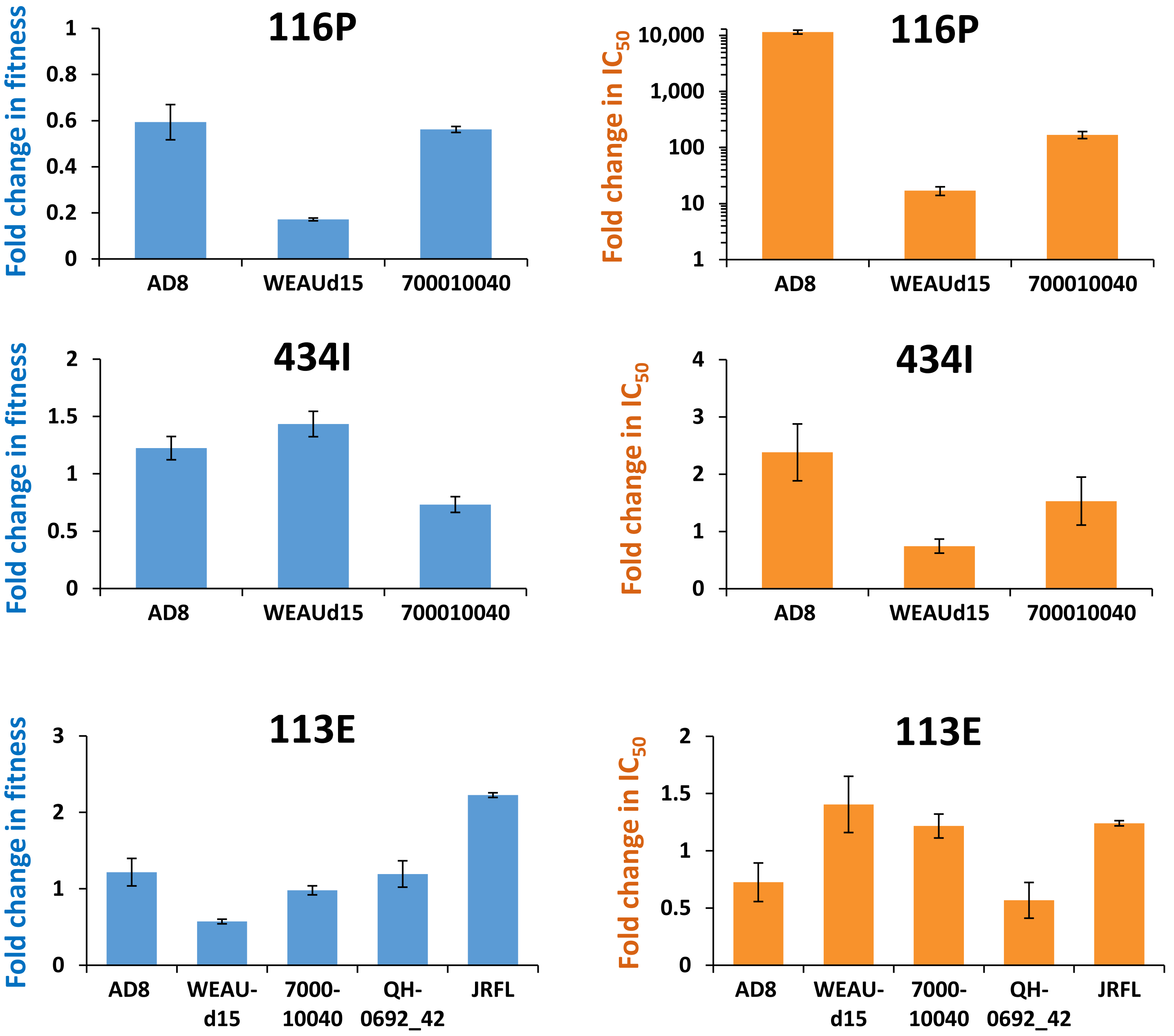


**Supplemental Figure S10. Effects of mutations on Env fitness and TMR resistance in diverse isolates.** The indicated mutations were introduced in AD8 Env and in the T/F strains WEAUd15.410.5017 (accession number EU289202) and 700010040.C9.4520 (EU576418). Effects on Env fitness and resistance to TMR are expressed as their fold change relative to the corresponding wild-type Envs. Given the high emergence frequency of 113E after treatment, we also examined the effects of this mutation on strains QH-0692_42 (AY835439) and JRFL (U63632). Error bars, standard errors of the means (SEM).

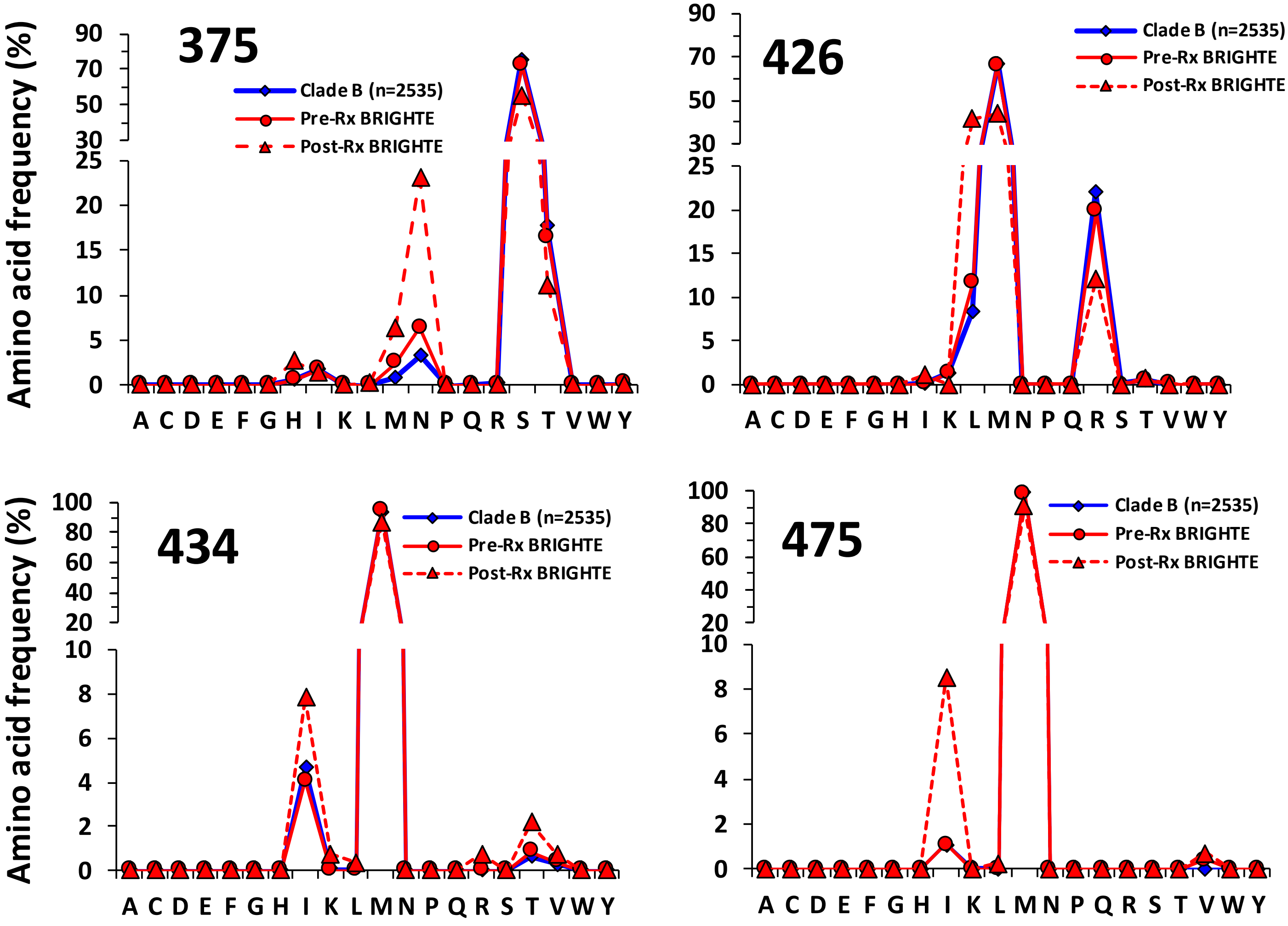


**Supplemental Figure S11.** Frequency distribution of all amino acids at the indicated positions of Env. Frequencies are expressed as a percent of all variants at that position and are shown for: **(i)** The population HIV-1 of clade B, represented by the panel of 2,535 Envs, **(ii)** Samples collected in the BRIGHTE trial before treatment, and **(iii)** Samples collected in the BRIGHTE trial after treatment.

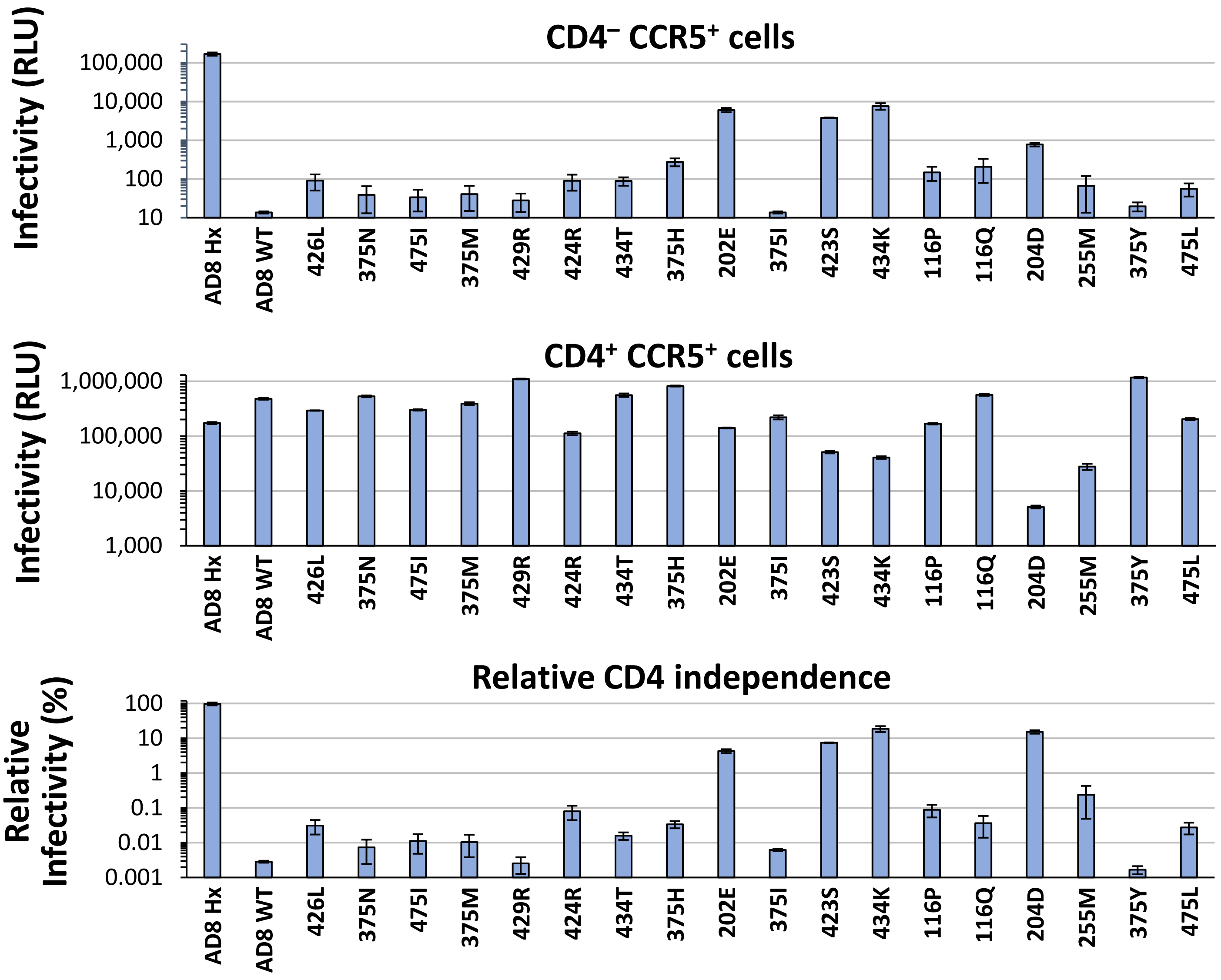


**Supplemental Figure S12. CD4-independent infection by REMs.** The indicated mutations were introduced in AD8 Env, and pseudoviruses were used to infect Cf2Th cells that stably express human CCR5 or both CD4 and CCR5. The relative CD4 independence is calculated as the level of infection measured in the CCR5+ cells as a percent of infection measured in the CD4+CCR5+ cells. The CD4-independent mutant AD8/Hx(N197S) was used as a positive control. REMs appear from left to right in decreasing order of frequency in the escape group. Error bars, SEM.

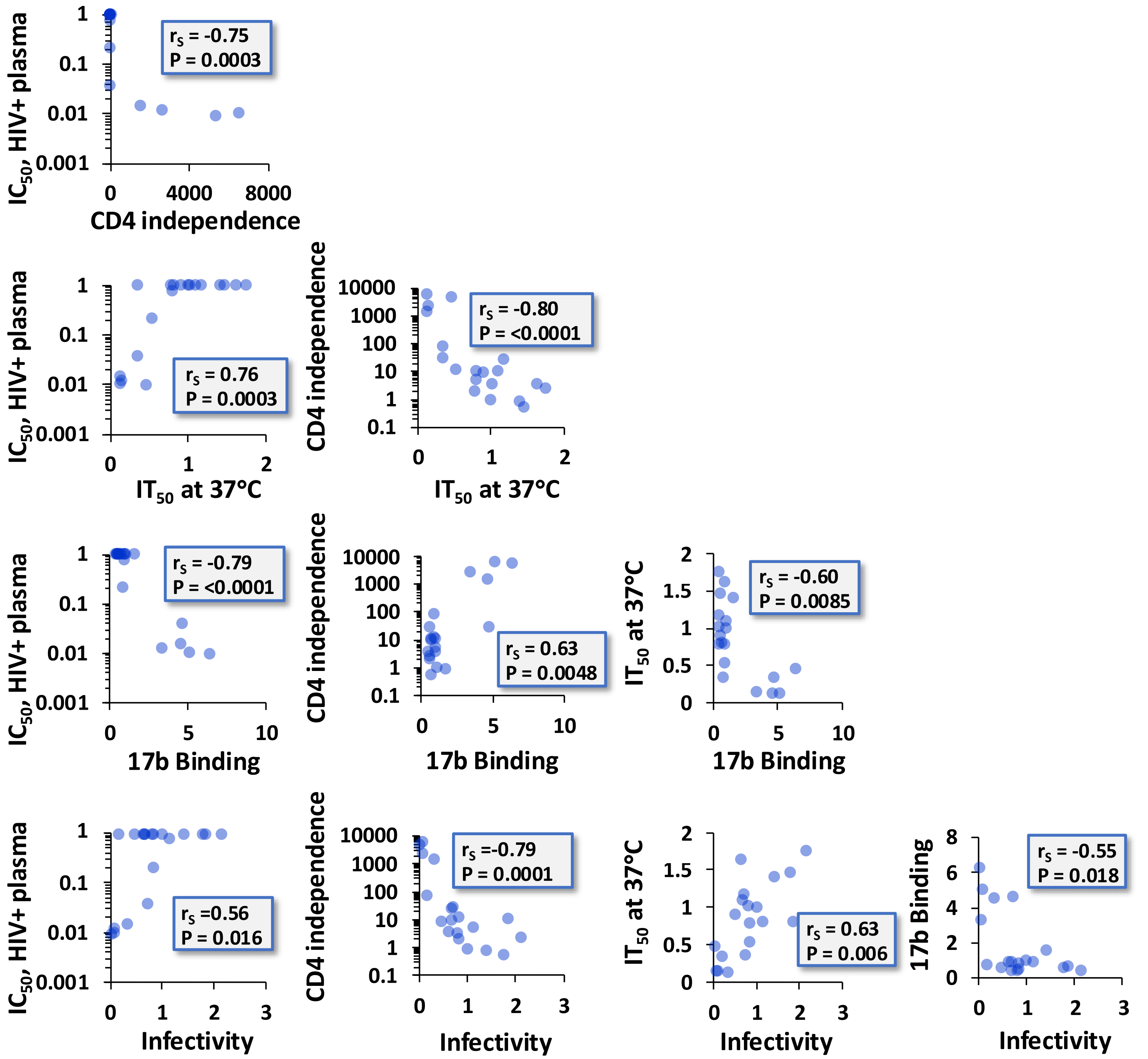


**Supplemental Figure S13.** Relationships between Env features we measured for the 18 REMs. All values describe the effects of the REMs relative to the wild-type AD8 Env. $r_S$, Spearman rank correlation coefficient; P-value, two-tailed test.

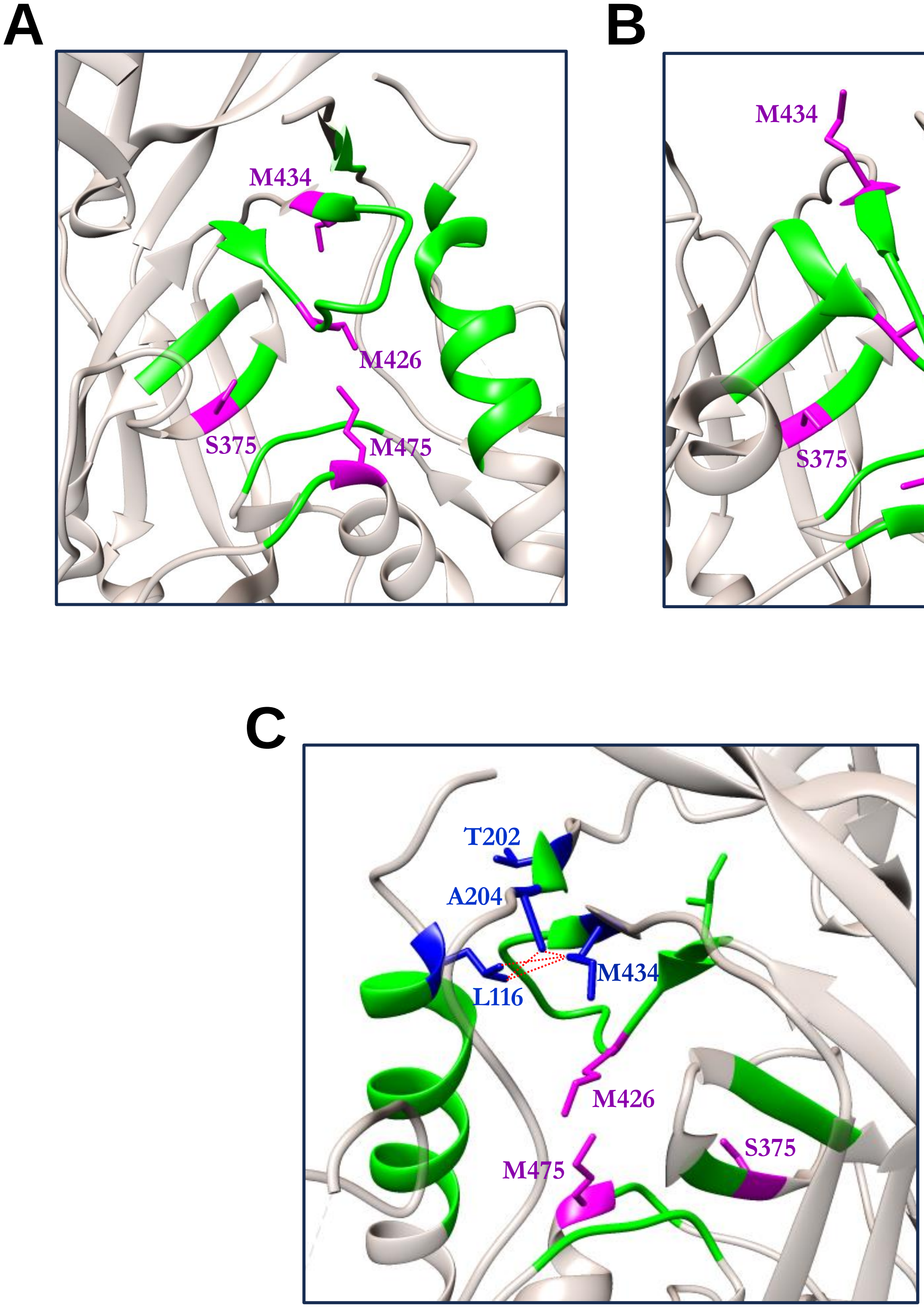


**Figure S14. Mechanism of Env resistance to TMR. (A,B)** Arrangement of the $SM^3$ sites (in pink) in the unliganded (A) and CD4-bound (B) forms of the clade B Env B41 (PDB IDs 6U59 and 5VN3, respectively). **(C)** Clasp-like structure that we propose maintains Env in the native state, formed by L116, M434 and A204. Mutations at these positions increase exposure of the CoR-BS, enhance sensitivity to non-neutralizing plasma from HIV-infected individuals, reduce Env fusion competence and increase virus inactivation rate at 37°C. The $SM^3$ sites 375, 426 and 475 are shown for orientation as well as position 202, which is also associated with transition to a CD4-bound-like conformation.

**A**

**From nucleotide** (columns) / **To nucleotide** (rows)

| | A | C | T | G |
|---|---|---|---|---|
| A | | 5.9 | 1.9 | 25.0 |
| C | 4.7 | | 22.6 | 1.7 |
| T | 8.2 | 7.4 | | 8.8 |
| G | 10.8 | 0.4 | 2.7 | |

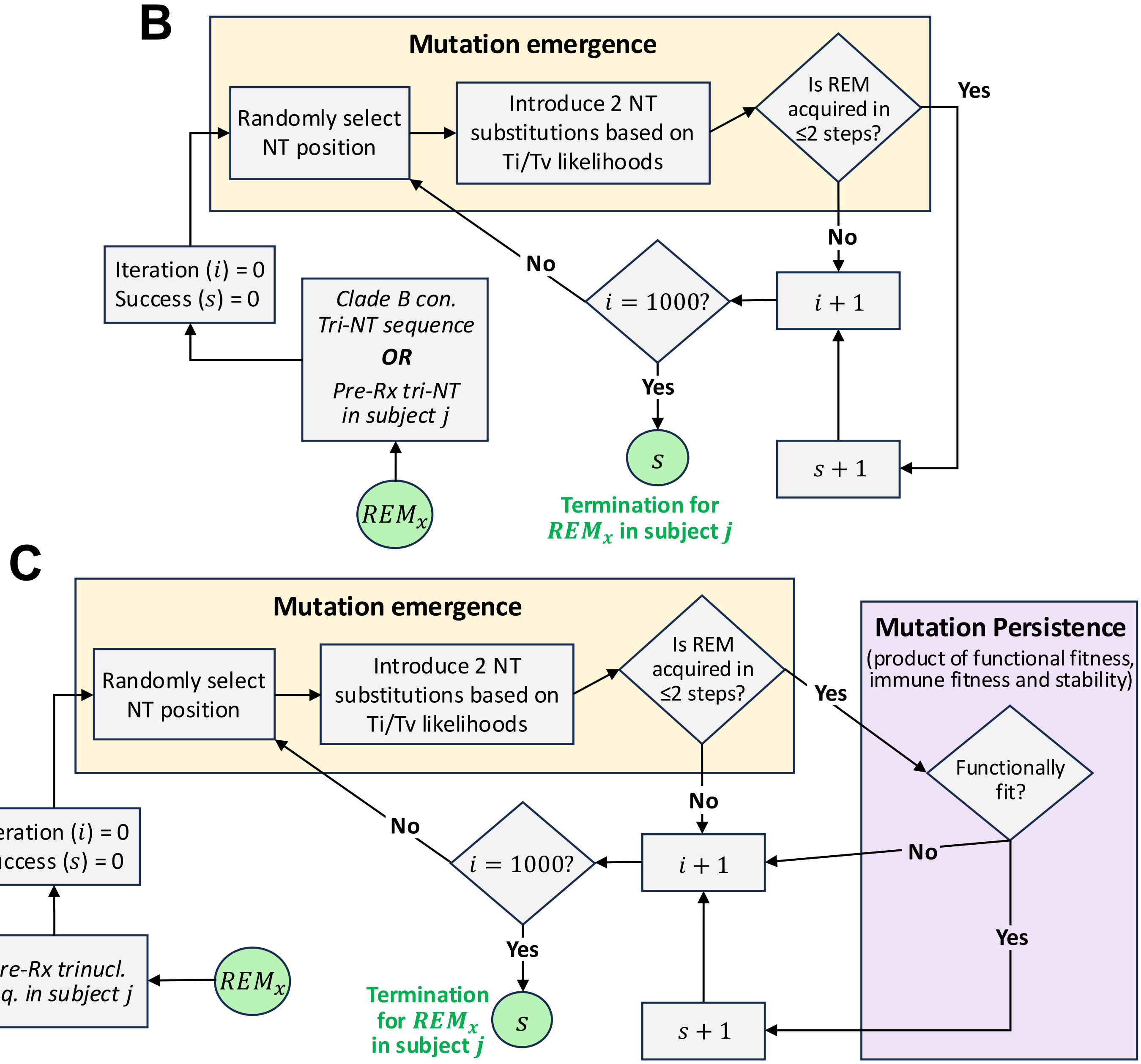


**Figure S15. Simulation of the emergence and persistence of REMs in the population and in TMR-treated individuals. (A)** Base substitution types detected in an acellular assay using reverse transcriptase of HIV-1 strain BH10, as described by Martínez del Río, *et al.*, 2024. Values are expressed as a percent of all base substitutions detected. **(B)** Simulation to determine the likelihood for appearance of REMs within two NT substitutions, using the clade B consensus sequence at each position or the pre-treatment sequences from the BRIGHTE participants. The number of success events ($s$) from the 1000 iterations ($i$) is calculated for each subject and averaged across the 65 subjects of the escape group. **(C)** Augmentation of the above algorithm with a decision module that accounts for the combined fitness of each REM.

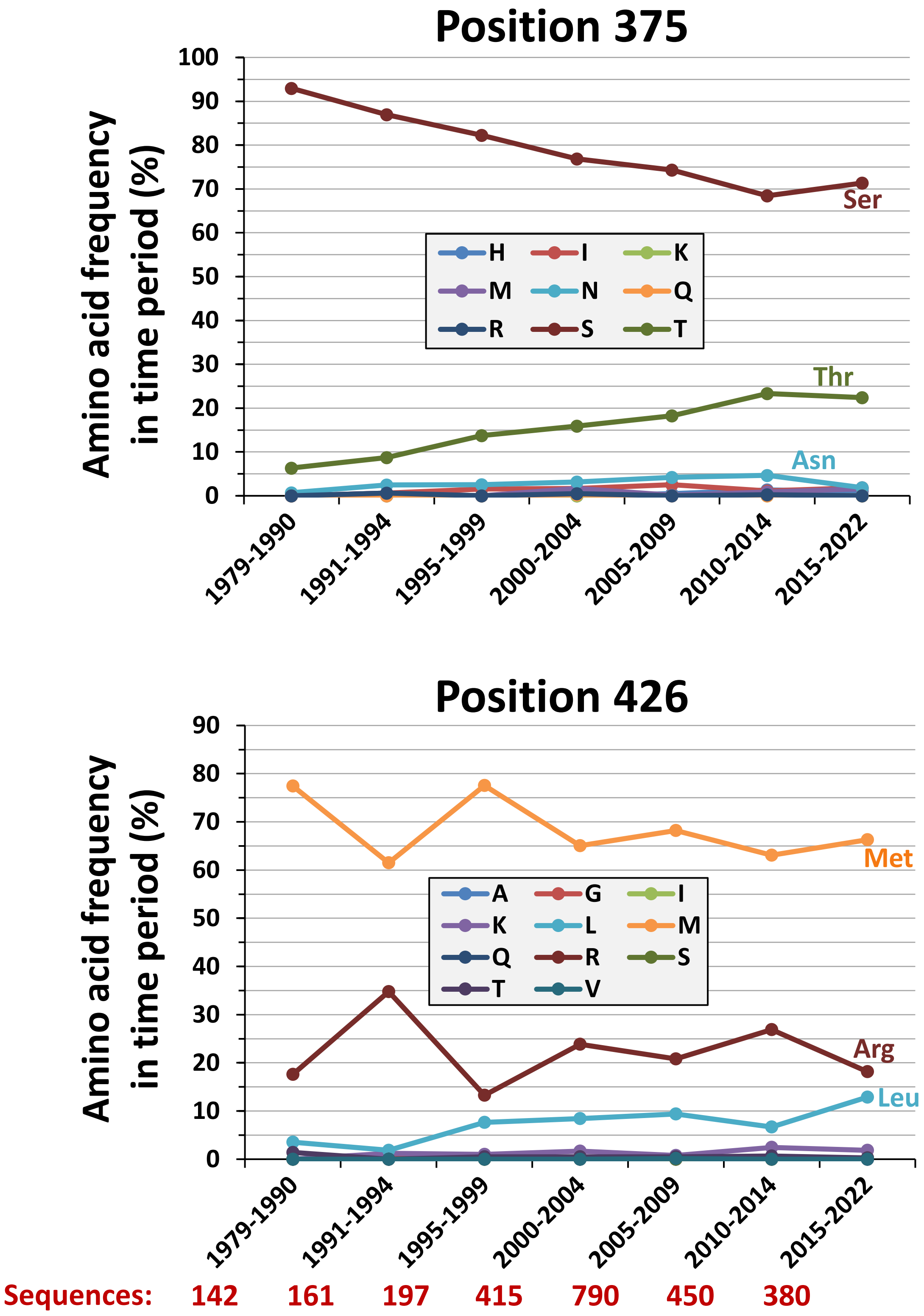


**Supplemental Figure S16. Historical changes in amino acid occupancy at positions 375 and 426 in HIV-1 clade B.** The 2535 sequences of the fostemsavir-untreated subjects infected by HIV-1 clade B were partitioned according to the year of sample collection. The frequency of all amino acids in each time period was calculated. The number of sequences in each time period is shown below the graphs in red.

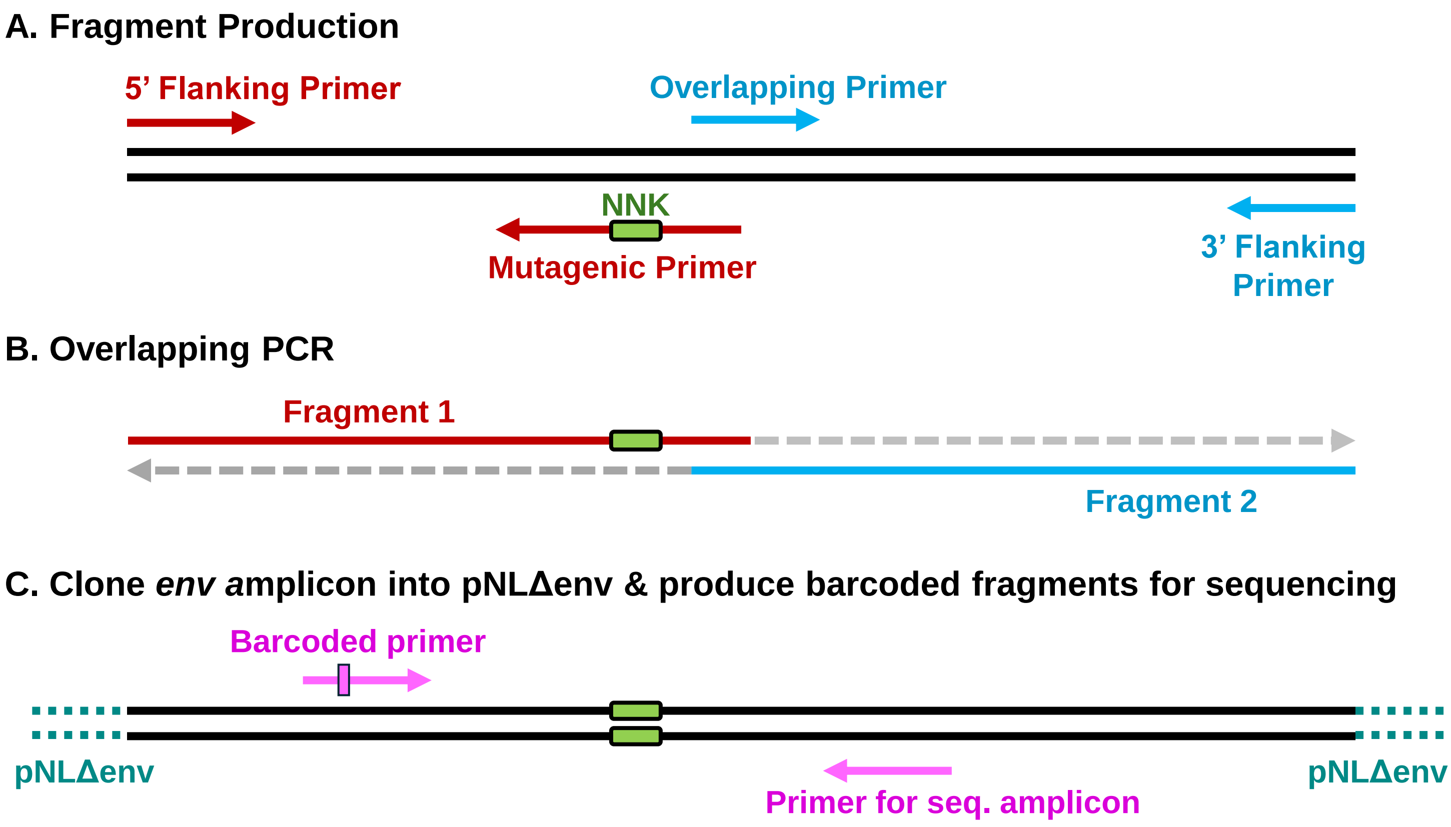


| Purpose | Primer | Sequence |
|---|---|---|
| **S375X Fragment 1** | 5' Flanking | TTTCATGACAAAAGCCTTAGGCATCTCC |
| **S375X Fragment 1** | Mutagenic | GAATTACAGTAGAAAAATTCGCCACCACAATTAAAMNNGTG CATTACAATTTCTGGGTCC |
| **S375X Fragment 2** | Overlapping | GTGGTGGCGAATTTTTCTACTGTAATTCAACAC |
| **S375X Fragment 2** | 3' Flanking | GTTTTTCTAGGTCCCGAGATACTGCTCC |
| **M426X Fragment 1** | 3' Flanking | GTTTTTCTAGGTCCCGAGATACTGCTCC |
| **M426X Fragment 1** | Mutagenic | CTATCACACTCCCCTGTAGAATTAAACAAATTATAAACNNKT GGCAAGAAGTAGGAAAAG |
| **M426X Fragment 2** | 5' Flanking | TTTCATGACAAAAGCCTTAGGCATCTCC |
| **M426X Fragment 2** | Overlapping | GTTTATAATTTGTTTAATTCTACAGGGGAGTGTGATAGTGTC |
| **Sequencing** | Barcoded Index 1 | CCAGTACTGTCAACTCAACTGCTGTTAAATGG |
| **Sequencing** | Barcoded index 2 | CCAGTATTGTCAACTCAACTGCTGTTAAATGG |
| **Sequencing** | Barcoded index 3 | CCAGTAATGTCAACTCAACTGCTGTTAAATGG |
| **Sequencing** | Rev sequencing | GTTGTTCTGCTGTTGCACTATACC |
| **pNLΔenv linearization** | pNLΔenv Fw | ACTGCTGCCTAGCGCTTTTGTCATGAAACAAAC |
| **pNLΔenv linearization** | pNLΔenv Rev | AGGCAGCAGTATCCCGGGACCTAGAAAAACATG |

**Supplementary Figure S17. Production of plasmid libraries for deep mutational scanning of positions 375 and 426.** Fragment 1 is produced by PCR using a universal 5′ flanking primer and a mutagenic primer containing the NNK codon. Fragment 2 is generated using an overlapping primer with 20–30 bp of homology downstream of the NNK site and a universal 3′ flanking primer. Fragments 1 and 2 are joined by overlap-extension PCR to yield the full-length amplicon that spans the *env* gene. The amplicon is then cloned into the env-deleted proviral vector pNL*Δ*env by the InFusion method. Barcoded primers were used to amplify the region containing the NNK site to allow multiplexing of samples in sequencing reactions.
