## Supplemental Tables S1-S3 for "Comprehensive Mapping of the Virus and Host Factors that Guide the Paths of HIV-1 Escape from a Therapeutic"

**Supplemental Table S1. Env positions used as input for the XGBoost model to estimate resistance to TMR.**

| Env Position[a] | Distance (Å)[b] | Env atom[c] | TMR atom[d] |
|---|---|---|---|
| **113** | 1.65 | OD1 | H23 |
| **255** | 2.33 | O | H3 |
| **432** | 2.51 | OE1 | H21 |
| **375** | 2.52 | OG | H6 |
| **427** | 2.70 | CE2 | H2 |
| **433** | 2.85 | O | H19 |
| **109** | 2.88 | O | H13 |
| **424** | 2.94 | CG1 | H18 |
| **425** | 3.00 | O | H17 |
| **112** | 3.02 | CZ2 | H9 |
| **202** | 3.09 | CG2 | H19 |
| **376** | 3.17 | N | H14 |
| **434** | 3.25 | CE | H18 |
| **475** | 3.41 | SD | H1 |
| **370** | 3.44 | OE1 | H5 |
| **384** | 3.50 | OH | H5 |
| **426** | 3.60 | N | H17 |
| **116** | 4.07 | CD1 | C29 |
| **256** | 4.19 | N | H3 |
| **108** | 4.45 | O | O09 |
| **428** | 4.68 | N | O03 |
| **383** | 4.71 | O | H6 |
| **257** | 4.73 | N | H14 |
| **382** | 4.85 | CZ | O06 |
| **110** | 4.94 | N | H13 |
| **200** | 5.00 | O | H19 |
| **377** | 5.05 | N | H3 |
| **117** | 5.34 | CA | H20 |
| **114** | 5.59 | N | H22 |
| **201** | 5.70 | C | H19 |
| **479** | 6.01 | CZ2 | H10 |
| **69** | 6.02 | CZ2 | H9 |
| **374** | 6.02 | C | H14 |
| **254** | 6.02 | C | H3 |
| **111** | 6.06 | N | H13 |
| **373** | 6.19 | O | H6 |
| **435** | 6.22 | N | H15 |
| **423** | 6.32 | C | H17 |
| **105** | 6.51 | O | O09 |
| **121** | 6.55 | CE | N32 |
| **253** | 6.60 | O | H9 |
| **371** | 6.68 | N | H6 |
| **431** | 6.69 | O | H19 |
| **369** | 6.70 | O | H6 |
| **473** | 6.80 | O | H1 |
| **478** | 6.87 | OD1 | H3 |
| **203** | 6.93 | N | H19 |
| **421** | 7.00 | CE | H5 |
| **429** | 7.03 | N | O03 |
| **210** | 7.16 | CG | O06 |
| **106** | 7.21 | O | H13 |
| **199** | 7.33 | OG | H19 |
| **115** | 7.34 | N | H22 |
| **422** | 7.49 | O | H18 |
| **107** | 7.49 | O | O09 |
| **474** | 7.49 | C | H1 |

| Env Position | Distance (Å) | Env atom | TMR atom |
|---|---|---|---|
| **368** | 7.59 | OD2 | H5 |
| **436** | 7.68 | N | H18 |
| **118** | 7.68 | N | H20 |
| **430** | 7.73 | O | N28 |
| **204** | 7.91 | N | H19 |
| **68** | 7.97 | CG2 | H7 |
| **258** | 7.99 | N | H14 |
| **259** | 8.17 | O | H14 |
| **261** | 8.26 | CD1 | H14 |
| **120** | 8.30 | O | H19 |
| **385** | 8.31 | N | H6 |
| **476** | 8.39 | N | H1 |
| **378** | 8.44 | N | H3 |
| **381** | 8.48 | O | C23 |
| **119** | 8.48 | O | H20 |
| **104** | 8.56 | O | O09 |
| **380** | 8.71 | O | O06 |
| **372** | 8.87 | N | H6 |
| **419** | 8.92 | O | H6 |
| **195** | 9.12 | OD1 | H19 |
| **122** | 9.13 | N | H19 |
| **212** | 9.20 | CD | O06 |
| **472** | 9.55 | CA | H14 |
| **211** | 9.61 | N | O06 |
| **420** | 9.83 | C | H6 |
| **209** | 9.86 | O | H9 |
| **471** | 9.93 | O | H14 |
| **453** | 10.13 | CD1 | H14 |
| **437** | 10.18 | CD | H18 |
| **205** | 10.20 | N | H19 |
| **180** | 10.24 | OD1 | H5 |
| **260** | 10.32 | N | H14 |
| **477** | 10.50 | N | H1 |
| **179** | 10.54 | CD1 | H6 |
| **123** | 10.56 | CG2 | N32 |
| **252** | 10.67 | O | H9 |
| **198** | 10.69 | OG1 | H15 |
| **67** | 10.79 | OD1 | H9 |
| **103** | 10.88 | O | O09 |
| **379** | 10.88 | N | O06 |
| **194** | 11.04 | CG2 | H17 |
| **387** | 11.09 | OG1 | H14 |
| **208** | 11.26 | CG2 | H8 |
| **418** | 11.38 | CB | H6 |
| **386** | 11.43 | N | H6 |
| **296** | 11.67 | SG | H14 |
| **331** | 11.72 | SG | H14 |
| **102** | 11.74 | O | O09 |
| **213** | 11.87 | N | H9 |
| **447** | 12.02 | OG | H3 |
| **101** | 12.08 | O | H10 |
| **445** | 12.09 | SG | H14 |
| **196** | 12.14 | N | H19 |
| **262** | 12.15 | ND2 | H3 |
| **294** | 12.38 | CD1 | H14 |
| **367** | 12.38 | C | H6 |
| **217** | 12.42 | OH | O09 |

[a] Positions located within 4Å, 5Å, 7.5Å, 10Å and 12.5Å are highlighted in different shades of blue.

[b] The minimal distance between each Env position and any TMR atom was calculated using coordinates of the TMR-bound structure (PDB ID 5U7O).

[c,d] The Env and TMR atoms involved in the interactions.

**Supplemental Table S2. Structure-guided approach to identify mutations that may increase Env resistance to TMR.**

| Position [a] | 109 | 112 | 113 | 202 | 255 | 375 | 424 | 426 | 427 | 432 | 434 | 116 | 257 | 370 | 382 | 384 | 475 |
|---|---|---|---|---|---|---|---|---|---|---|---|---|---|---|---|---|---|
| Distance from TMR [b] | 2.9 | 3.0 | 1.7 | 3.1 | 2.3 | 2.5 | 2.9 | 3.6 | 2.7 | 2.5 | 3.3 | 4.1 | 4.7 | 3.4 | 4.9 | 3.5 | 3.4 |
| Interacts with TMR? [c] | Yes | Yes | Yes | Yes | Yes | Yes | Yes | Yes | Yes | Yes | Yes | No | No | No | No | No | No |
| Ala [d] | 0 | 0 | 0.1 | 0.1 | 0.6 | 0 | 0.04 | 0.04 | 0 | 0 | 0 | 0 | 0.04 | 0.1 | 0 | 0 | 0 |
| Arg | 0 | 0 | 0 | 1.2 | 0 | 0.2 | 0.04 | 22.1 | 0.2 | 7.4 | 0 | 0 | 0 | 0 | 0 | 0 | 0 |
| Asn | 0 | 0 | 0.1 | 0.04 | 0 | 3.3 | 0 | 0 | 0 | 0.1 | 0 | 0 | 0.04 | 0.04 | 0 | 0 | 0 |
| Asp | 0 | 0 | 98.2 | 0 | 0 | 0 | 0 | 0 | 0 | 0 | 0 | 0 | 0 | 0.04 | 0 | 0 | 0 |
| Cys | 0 | 0 | 0 | 0 | 0 | 0 | 0 | 0 | 0.12 | 0 | 0 | 0 | 0 | 0 | 0 | 0 | 0 |
| Glu | 0 | 0 | 1.7 | 0 | 0 | 0 | 0 | 0 | 0 | 0.1 | 0 | 0.04 | 0 | 99.8 | 0 | 0 | 0 |
| Gln | 0 | 0 | 0 | 0.04 | 0 | 0.04 | 0 | 0.04 | 0 | 0.7 | 0 | 0 | 0 | 0.04 | 0 | 0 | 0 |
| Gly | 0 | 0.04 | 0 | 0 | 0 | 0 | 0 | 0.04 | 0.12 | 0 | 0 | 0 | 0 | 0 | 0 | 0 | 0 |
| His | 0 | 0 | 0 | 0 | 0 | 0.6 | 0 | 0 | 0 | 0 | 0 | 0 | 0 | 0 | 0 | 0 | 0 |
| Ile | 98.3 | 0 | 0 | 0.04 | 2.0 | 1.7 | 78.8 | 0.08 | 0 | 0.1 | 4.7 | 0.3 | 0.04 | 0 | 0 | 0 | 1.1 |
| Leu | 0.04 | 0 | 0 | 0 | 0.04 | 0 | 0 | 8.3 | 0 | 0 | 0.1 | 99.6 | 0.04 | 0 | 0 | 0 | 0.04 |
| Lys | 0.4 | 0 | 0 | 4.5 | 0 | 0.04 | 0 | 1.4 | 0 | 91.2 | 0.1 | 0 | 0 | 0.04 | 0 | 0 | 0.04 |
| Met | 0 | 0 | 0 | 0 | 0.1 | 0.8 | 0.04 | 67.4 | 0 | 0 | 94.2 | 0.04 | 0 | 0 | 0 | 0 | 98.8 |
| Phe | 0 | 0 | 0 | 0 | 0 | 0 | 0 | 0 | 0 | 0 | 0 | 0.04 | 0 | 0 | 99.9 | 2.1 | 0 |
| Pro | 0 | 0 | 0 | 0 | 0 | 0 | 0 | 0 | 0 | 0 | 0 | 0 | 0.1 | 0 | 0 | 0 | 0 |
| Ser | 0.1 | 0 | 0 | 0.2 | 0.04 | 75.7 | 0 | 0.1 | 0 | 0 | 0 | 0 | 0.04 | 0 | 0 | 0 | 0 |
| Thr | 0.2 | 0 | 0 | 94.0 | 0 | 17.7 | 0 | 0.5 | 0 | 0.4 | 0.6 | 0 | 99.7 | 0 | 0 | 0 | 0 |
| Trp | 0 | 100.0 | 0 | 0 | 0 | 0 | 0 | 0 | 99.6 | 0 | 0 | 0 | 0 | 0 | 0 | 0 | 0 |
| Tyr | 0 | 0 | 0 | 0 | 0 | 0 | 0.04 | 0 | 0 | 0 | 0 | 0 | 0 | 0 | 0.1 | 98.0 | 0 |
| Val | 1.0 | 0 | 0 | 0 | 97.3 | 0 | 21.1 | 0.04 | 0 | 0.04 | 0.3 | 0.04 | 0 | 0 | 0 | 0 | 0 |

[a] Env positions with side chains that occupy the CD4-binding pocket and are within 5Å of the TMR molecule were identified using the TMR-liganded structure of Env (PDB ID 5U7O).

[b] Distance (in Å) between the closest atoms of the TMR molecule and the residue found at the indicated position on the structure.

[c] Absence or presence of an interaction between the side chain of the indicated amino acid and TMR, as suggested by the above structure.

[d] The frequency of all amino acids at the indicated positions was calculated using a panel of 2,535 isolates from HIV-1 clade B. Residues with a frequency higher than 0.5% at each site were further analyzed for their effects on Env fitness and resistance to TMR.

**Supplemental Table S3. Summary of all mutations identified by the four approaches as suspected of increasing Env resistance to TMR.**

| Position | AA in con. B | GB Regressor [a] | Probabilistic model[b] | Structure guided [c,d] | Previously published | |
|---|---|---|---|---|---|---|
| | | | | | Mutation [e] | Reference |
| 109 | Ile | | | Val | | |
| 113 | Asp | Glu | Ala, Glu, His | Glu, Gly | | |
| 116 | Leu | | | | Pro, Gln | Zhou et al, 2014 |
| 200 | Val | | | Ser | | |
| 202 | Thr | | | Ala, Lys, Ser, Arg | Glu | Gartland et al., 2024 |
| 204 | Ala | | | | Asp | Zhou et al, 2014 |
| 210 | Phe | Trp | | | | |
| 255 | Val | Ile | Ala | Ala, Ile, Met | | |
| 373 | Met | Thr | Gln | | | |
| 375 | Ser | Thr, His, Asn | His, Met, Tyr, Ile | Asn, His, Ile, Met, Thr | His, Met, Thr | Pancera et al, 2017, Zhou et al, 2014 |
| 376 | Phe | | Ile, Leu | | | |
| 377 | Asn | | Val | | | |
| 384 | Tyr | | Leu | Phe, Cys | | |
| 423 | Ile | | Tyr | | Tyr, Phe, Ser | Zhou et al, 2010 |
| 424 | Ile | Val | Arg | Val | | |
| 426 | Met | Leu | Ile | Arg, Leu, Lys, Thr | Leu | Zhou et al, 2014, Ray et al, 2013 |
| 429 | Glu | Arg | His | | | |
| 432 | Lys | Gln | Leu | Arg, Gln | Leu | Pancera et al, 2017 |
| 434 | Met | Lys, Ile | | Ile, Thr, Leu | Ile, Lys | Zhou et al, 2014, Lataillade et al, 2018 |
| 475 | Met | Ile | | Ile, Leu | Ile | Pancera et al, 2017; Zhou et al, 2014 |
| 478 | Asn | | Ile | | | |
| 506 | Val | | | | Met | Ray et al, 2013 |
| 595 | Ile | | | | Phe, Val | Zhou et al, 2010 |
| 655 | Lys | | | | Glu, Ala | Zhou et al, 2010 |

[a] The 14 mutations with the highest SHAP values are indicated.

[b] Mutations identified by the probabilistic model.

[c] All Env positions with side chains that occupy the CD4-binding pocket and are within 5Å of the TMR molecule were analyzed for the frequency of each amino acid in the panel of 2,535 clade B Envs. All residues with a frequency of 0.5% or higher are shown. See all data in Table S2.

[d] Mutations in red font were selected based on the structure, but their prevalence in the population was lower than 0.5%.

[e] Mutations in red font were not explicitly noted in the references but were added due to suspected effects.
